# A Droplet Digital PCR Assay for Quantification of Bacteriophage Viral Vector Titer and Purity

**DOI:** 10.1101/2024.11.20.624577

**Authors:** Peter J. Voorhees, Ronni M. Ponek, Jamie D. Liu, Samuel K. Lai

## Abstract

**Purpose:** Bacteriophage (phage) based vectors offer considerable promise as tools for tuning the microbiome with molecular and genetic precision. However, standardized methods to rigorously characterize phage vectors remain lacking. Here, we present an optimized digital droplet PCR (ddPCR)-based assay for quantifying both the purity and potency of phage vector preparations.

**Methods:** We utilized central composite design to develop a ddPCR assay capable of quantifying the number of phage vector capsids packed with the phage vector genome or packed with the transgenic DNA of interest. This assay targets 2 unique DNA barcodes, designed to be biologically inert and maximally orthogonal to existing DNA sequences.

**Results:** Through stringent optimization, we were able to achieve assay conditions that enable a dynamic range of nearly 3 orders of magnitude and correct for systemic error in the assay. We then show that biological activity assays consistently underestimate transgene-packed vectors titers, leading to overestimation of true transduction efficiency, particularly when contamination by genome-packed vectors is high. We further demonstrate how this approach facilitates optimization of vector production conditions and substantially improves the precision and reproducibility of phage vector transduction.

**Conclusion:** Compared to assays of biological activity, this optimized ddPCR assay has improved accuracy and, through design of experiments optimization, high precision (CVs = 5.5 ± 1.3% and 4.5 ± 1.0% for the genome and transgene barcodes, respectively). This assay can be broadly adopted to characterize and quality control vector preparations for various applications.

## Introduction

Bacteriophages (phages), viruses that infect bacteria, have long been studied for their potential as antimicrobial agents [1–3]. Recently, there is growing interest in their use as vehicles to deliver transgenic DNA that modulates the function of target bacteria, without inducing lysis of transduced bacteria [4–7]. For instance, engineered phage vectors can suppress shiga toxin production in susceptible *Escherichia coli* (*E. coli*) in the mammalian gut, thereby ameliorating the pathogenic effects of these strains without disrupting an ecological niche that other opportunistic pathogens might colonize [8–10]. The ability of phage vectors to precisely perform genetic manipulations on individual microbiome constituents *in situ* underscores their value as an exciting new approach in microbiome engineering.

To support further translational development, assays that offer robust and accurate characterization of phage vectors must be developed. In human gene therapy, there are established quality control and quantification assays to assess safety, potency, purity and stability of viral vectors, all of which are critical to their safe and efficacious use in clinical trials. Indeed, accurate and reproducible quantitation of transduction potency is essential for consistent dosing, which in turns involves measuring the precise amount (titer) of a viral vector in a given volume, including both the amount of transgenic DNA present as well as the number of biologically active particles [11]. Transgene copies in viral vectors for human gene therapy are often quantified by either qPCR or digital droplet PCR (ddPCR) [12,13], whereas transgene expression in permissive cell lines is used to determine the number of biologically active particles—those capable of mediating successful delivery of transgenic DNA (i.e. transduction) [14,15]. Measures of purity must account for residual proteins and nucleic acids from the vector production cell line, helper viruses or plasmids [16], as well as empty vectors and vector capsids packed with non-target DNA, including viral genomic DNA.

While the importance of these quality control measures in developing effective and reproducible treatments have been fully established in viral vectors for human gene therapy, analogous assays that determine potency and purity for phage vectors designed to genetically reprogram the microbiome have not been established. Similar to mammalian viral vectors, phage vector transduction efficiencies can be affected by numerous factors, including the presence of cofactors, and the size, complexity, and stability of the packed transgenic DNA, and thus can vary widely. As a result, standard biological activity assays are unlikely to accurately reflect the number of transgene-packed phage vectors. This is particularly important for phage vectors generated by modifying lytic phages, as the bacterial cytotoxicity of genome-packed vectors can significantly impact measured transduction levels due to target cell killing [17,18]. We believe quantitative PCR assays are therefore essential in not only understanding readouts of biological activity assays but also informing the appropriate conditions with which to run these assays.

To meet this need, we report here a ddPCR-based assay for quantifying both the potency and purity of phage vectors by measuring the quantity of capsids packed with transgenic DNA and vector genome DNA. This assay was first optimized using central composite design (CCD), then compared against methods for measuring biologically active vector particles derived from both lytic and temperate phages. Finally, we demonstrate how this assay plays an essential role in optimizing the production of high-titer phage vectors and in ensuring accuracy and reproducible transduction across distinct vector preparations and replicates. As this assay is based on small, standardized DNA barcodes, it is readily applicable to a wide variety of phage vectors and we anticipate it will be crucial in the development of effective and reproducible phage vector products.

## Materials and Methods

### Barcode generation

A python script (DOI: 10.5281/zenodo.10869457) was written to computationally design the DNA barcode sequences. Briefly, this script generates a specified number of random DNA sequences of a set length, with a G/C content between 40-60%, without homopolymeric repeats >4 bp, and without common restriction enzyme sites, common promoter and RBS consensus sequences. The secondary structure of each sequence is then analyzed by RNAfold along a sliding 50 bp window across each entire sequence. The maximum minimum free energy (MFE) and average MFE are calculated for each sequence and sequences with a max MFE greater than a specified threshold are removed. The remaining sequences are sorted by average MFE, and bottom 90% are removed. Remaining sequences are then run through a BLAST search against the NCBI Nucleotide database and the percent identity, alignment length, E-value, Bit Score, and Query Coverage of the top BLAST match for each barcode is recorded. Next, the script finds the best barcodes based on minimal similarity to existing sequences, as determined by the BLAST search results, and minimal predicted secondary structure. Finally, the script finds the best pairs of barcodes. Ranking each pair by various criteria including max MFE, average MFE, alignment length, percent identity, and query coverage. The pairs are then sorted by weighted average rank and intra pair orthogonality (Hamming distance), and the results are saved to a csv file.

### Plasmid cloning

8673 and 7898 barcode reference plasmids (pBc8673 and pBc7898) were cloned by Twist Bioscience into pTwist Amp High Copy backbones. The transgene phagemids (pT7kan-mScarlet and pP1kan-mScarlet) and gp17 helper plasmid (pT7gp17) were cloned through Gibson HiFi (New England Biolabs, cat. no. E2621L) assembly of synthesized DNA fragments (IDT) (sequences in **Supplementary Table 8**) according to **Supplementary Table 9**.

### Phage genome assembly & engineering

The barcoded T7 genome was generated through a previously described yeast-based assembly and rebooting protocol [19]. Briefly, the T7 genome was PCR amplified in fragments, excluding the gp17 coding sequence. A yeast artificial chromosome was also amplified with regions of homology to the termini of the T7 genome (primer sequences in **Supplementary Table 10**). A double stranded DNA fragment encoding the 7898 barcode flanked by 60 bp sequences homologous to the regions up and downstream of gp17 was synthesized by Twist Bioscience. 1-4 μg of each fragment was then transformed into EBY100 *S. cerevisiae* via electroporation. YAC-captured T7 genomic DNA was purified from the yeast and transformed into *E. coli* NEB5α containing a T7 gp17 trans-complementation plasmid. These *E. coli* were mixed with soft agar and plated on LB agar plates. Plaques were then picked to isolate gp17-deficient barcoded T7 phage. To remove the *pacAB* locus from the P1 genome, λRED-mediated recombineering was used to first remove IS1 from the P1 episome in an *E. coli* mod749::IS5 c1.100 P1 lysogen (ATCC BAA-1001). The *pacAB* locus was then replaced with a synthesized DNA fragment (Twist Bioscience) encoding the 7898 barcode and a chloramphenicol resistance cassette (**Supplementary Table 10**: PJV1345).

### ddPCR central composite design

Primers and Affinity Plus probes specific to the 8673 and 7898 barcodes were synthesized by IDT (**Supplementary Table 11**). To optimize the ddPCR assay a central composite design experiment was designed using JMP 17.0.0. The factor setting ranges described in **Supplementary Table 3** were used for the annealing temperature, primer concentration, probe concentration, ramp rate, and cycle number. The following four optimization goals were set: minimize 8673 relative error, minimize 7898 relative error, maximize 8673 droplet separation, and maximize 7898 droplet separation. Relative error was defined as the relative difference between the measured barcode copies/well and the actual barcode copies/well. Droplet separation was defined as the difference between the mean amplitude of the positive and negative droplets. ddPCR was performed with each of the factor combinations generated by JMP (**Supplementary Table 4**) according to the manufacturers recommended protocol (Bio Rad #1863024) using a Bio Rad QX200 automated droplet generator and droplet generation oil for probes (Bio Rad #1864110), a Bio Rad T100 thermal cycler, and a Bio Rad QX200 droplet reader. Data analysis and thresholding was performed using QX Manager, Standard Edition (Version 1.2). Relative errors and droplet separation were then input into a JMP Fit Least Squares model and the resulting desirability function was maximized.

### ddPCR performance metric characterization

5-fold serial dilutions of the barcode reference plasmids were made, ranging from 1.28×10^6^ to 16 molecule/μL. Input plasmid concentrations were calculated using the barcode plasmid molecular weight and the plasmid mass, as quantified by Qubit (Invitrogen #Q33230). ddPCR was performed on these plasmid dilutions using the CCD-derived settings (T_a_ = 63°C, primer concentration = 5.1 μM, probe concentration = 2.25 μM, ramp rate of 1°C/s, 50 cycles). The relative error and coefficients of variance were calculated for both barcode reference plasmids at each dilution point. Measured *vs.* actual values were plotted in log-log form and linear regression was performed (GraphPad Prism 10.1.1) on the range of sample points that appeared linear. The LLOD was calculated as the sum of the mean of the concentrations in the no template control wells and 3 times the standard deviation of these wells.

### T7 phage vector production

*E. coli* NEB5α was transformed with pT7kan-mScarlet and pT7gp17. These cells were cultured for 16 hrs overnight. Overnight cultures were passaged 1:100 into fresh media and cultured until an OD_600_ of 0.6 was reached. Unless otherwise specified, these cultures were spiked with 1×10^8^ gp17-deficient barcoded T7 helper phage. Infection was allowed to proceed for 4 hrs at 37°C with shaking. Infections were then centrifuged at 10,000 x G for 10 min and supernatant was 0.2 μm filtered to isolate phage vectors. For **Fig 5** experiments, a full factorial production run was designed using JMP 17.0.0. The factor setting ranges described in **Supplementary Table 6** were used for the production bacteria concentration, helper phage inoculum concentration, and the incubation time. The optimization goals were set as: maximize transgene-packed capsid concentration and minimize genome-packed capsid concentration.

### P1 phage vector production

The P1 mod749::IS5 c1.100 *E. coli* lysogen engineered to contain the 7898 barcode in place of the *pacAB* locus was transformed with pP1kan-mScarlet. These cells were then grown at 30°C for 16 hrs overnight. Overnight cultures were passaged 1:100 into fresh media and cultured at 30°C until an OD_600_ of 0.6 was reached. Cultures were then incubated at 42°C for 2 hours to induce P1 production. These cultures were then centrifuged at 10,000 x G for 10 min and supernatant was 0.45 μm filtered to isolate phage vectors.

### Phage DNA purification and ddPCR

Phage DNA was purified using the Norgen Phage DNA Isolate Kit (Norgen Biotek Corp. # 46800) following the manufacturers recommended protocol. Purified DNA was digested with Sca-I (NEB # r3122) and 3 5-fold dilutions were made of the completed digestion reaction. Each of these dilutions was analyzed by the optimized ddPCR reaction and the highest concertation in the dynamic range of the assay was used to calculate transgene and genome copies/mL.

### Colony formation transduction assays

*E. coli* NEB5α were cultured for 16 hrs overnight. Overnight cultures were passaged 1:100 into fresh media and cultures until an OD_600_ of 0.6 was reached. These cells were then mixed with 500 μL of each phage vector to a final concentration of 1×10^8^ cells/mL in a final volume of 1 mL. These transduction reactions were incubated at 37°C for 90 min with shaking. Cells were then pelleted at 16,000 x G for 5 min and resuspended in fresh LB. 4-fold dilutions of the transduced cells were made and plated on LB plates supplemented with kanamycin (50 μg/mL). Plates were incubated for 16 hrs at 37°C and colonies were counted to calculate the number of transduced cells (transductants) per mL. For **Fig 4** experiments, five 5-fold dilutions of T7 phage vectors were made prior to incubation with target cells. For MOI normalized transduction reactions, a volume of vectors required to achieve the specified MOI was mixed with LB media to achieve a final volume of 500 μL, prior to incubation with target cells.

### Trans-complementation plaque formation assays

*E. coli* NEB5α transformed with pT7gp17 were cultured for 16 hrs overnight. Overnight cultures were passaged 1:100 into fresh media and cultures until an OD_600_ of 0.6 was reached. 800 μL of these cells were mixed with 8 mL of molten soft agar and overlaid on prewarmed LB agar plates supplemented with ampicillin (100 μg/mL). 12 10-fold dilutions of each of the 5 5-fold phage vector dilutions were made and spotted onto the pT7gp17-containing NEB5α lawn plates. Plates were incubated for 16 hrs at 37°C and plaques were counted to calculate the number of plaque forming units (PFU) per mL. PFU/mL concentrations were plotted against ddPCR-determined genome copies/mL and 2-way ANOVA with Šídák’s multiple comparisons test was used to evaluate differences between the two measures at each of the 5 phage vector dilutions.

## Results

### Development of orthogonal DNA barcodes

ddPCR reactions commonly target amplification of a 60-200 bp DNA sequence [20]. To maximize modularity and transferability of the assay, we inserted unique 300 bp DNA barcodes into both the transgenic DNA intended for phage vector packing as well as the genome of the helper phage used to generate the vectors. Upon purification of encapsidated DNA from these vectors, this allows for the use of duplex ddPCR to probe for each barcode and quantify the concentration of transgene and genome-packed phage particles, in units of encapsidated DNA copies/mL (**Fig 1**).

**Fig 1.**
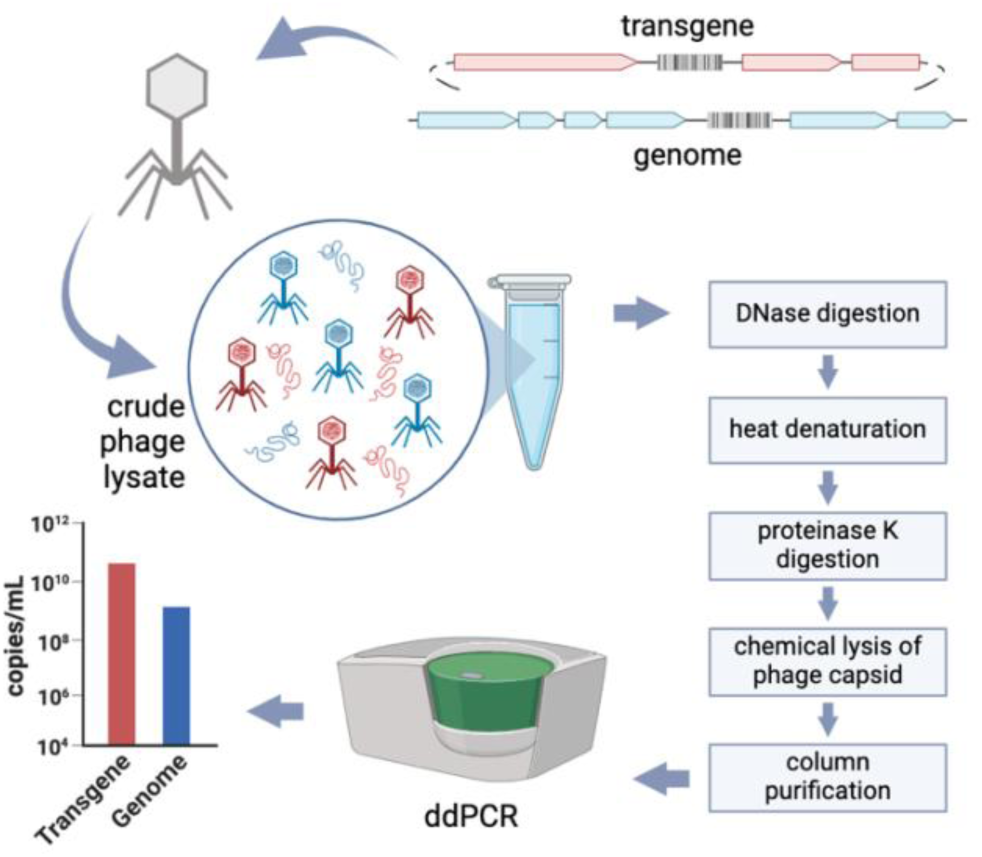
Phage vector characterization workflow using ddPCR. Unique DNA barcodes are inserted into the transgene cassette and phage genome. Both DNA are then used to produce phage vectors, which typically consists of mixed population packed with either transgene or the genome DNA. Following a purification protocol that digests non-encapsidated DNA, phage packed-DNA is recovered and analyzed via ddPCR to determine the absolute quantity and proportions of transgene and genome-packed vector particles

To choose DNA barcode sequences with high ddPCR compatibility and minimal similarity to naturally occurring DNA sequences, we focused on generating a pool of sequences with a G/C content between 40-60% and homopolymeric repeats <4 bps. We then filtered the barcode pool to remove those with common restriction endonuclease sites, consensus promoter or ribosome binding site sequences, and significant secondary structure. To maximize transferability to other vector systems, we then ranked the filtered pool to minimize similarity to sequences in the NCBI nucleotide database. Of those with the lowest similarity to reported DNA sequences, pairs of remaining barcodes were ranked for orthogonality to each other. From an initial pool of 10,000 candidates, we selected barcodes 8673 and 7898 for further development (**Supplementary Tables 1 and 2**).

### Central composite design identifies cycle number, annealing temperature, probe concentration, and ramp rate as key determinants of assay accuracy and droplet separation

Due to the exceptional sensitivity of ddPCR, reaction optimization is required to ensure accurate and reproducible readouts. The annealing temperature (T_a_), primer concentration, probe concentration, ramp rate, and number of PCR cycles can all influence the readout of a ddPCR assay. We therefore decided to optimize the duplex ddPCR assay using CCD (**Supplementary Tables 3 and 4**), with the goals of minimizing the relative error of the assay and maximizing the separation between positive and negative droplet populations for each barcode. To determine relative error, we used the assay to determine the concentrations of two reference plasmids, each encoding a single barcode (either 8673 or 7898). We also calculated the theoretical concentration of these barcode plasmids (copies/mL), which served as references for comparison to the measured barcode concentrations.

Applying a least-squares model to the data (**Supplementary Table 5**), we found a moderate fit for both 8673 and 7898 relative errors (R^2^ = 0.93 and 0.83, respectively) (**Figs 2A and 2B**), and a strong fit for both 8673 and 7898 droplet separation (R^2^ = 0.96 and 0.98, respectively) (**Figs 2C and 2D**). In this model, the most significant main effects were the number of cycles (p = 3.89×10^-6^), followed by the annealing temperature (p = 9.86×10^-4^), the probe concentration (p = 1.01×10^-3^), and the ramp rate (p = 1.30×10^-3^). Interactions between the ramp rate and cycles (p = 2.25×10^-3^) and the probe concentration and cycles (p = 9.95×10^-3^) were also significant (**Fig 2E**). The response surface graphs for relative error (**Fig 2F**) and separation (**Fig 2G**) as a function of ramp rate by cycles, and for relative error (**Fig 2F**) and separation (**Fig 2G**) as a function probe concentration by cycles appeared to indicate that the significance of these effects are primarily driven by the droplet separation. Notably, these data indicate a cycle number near 40 appears to minimize relative error for both 8673 and 7898.

**Fig 2.**
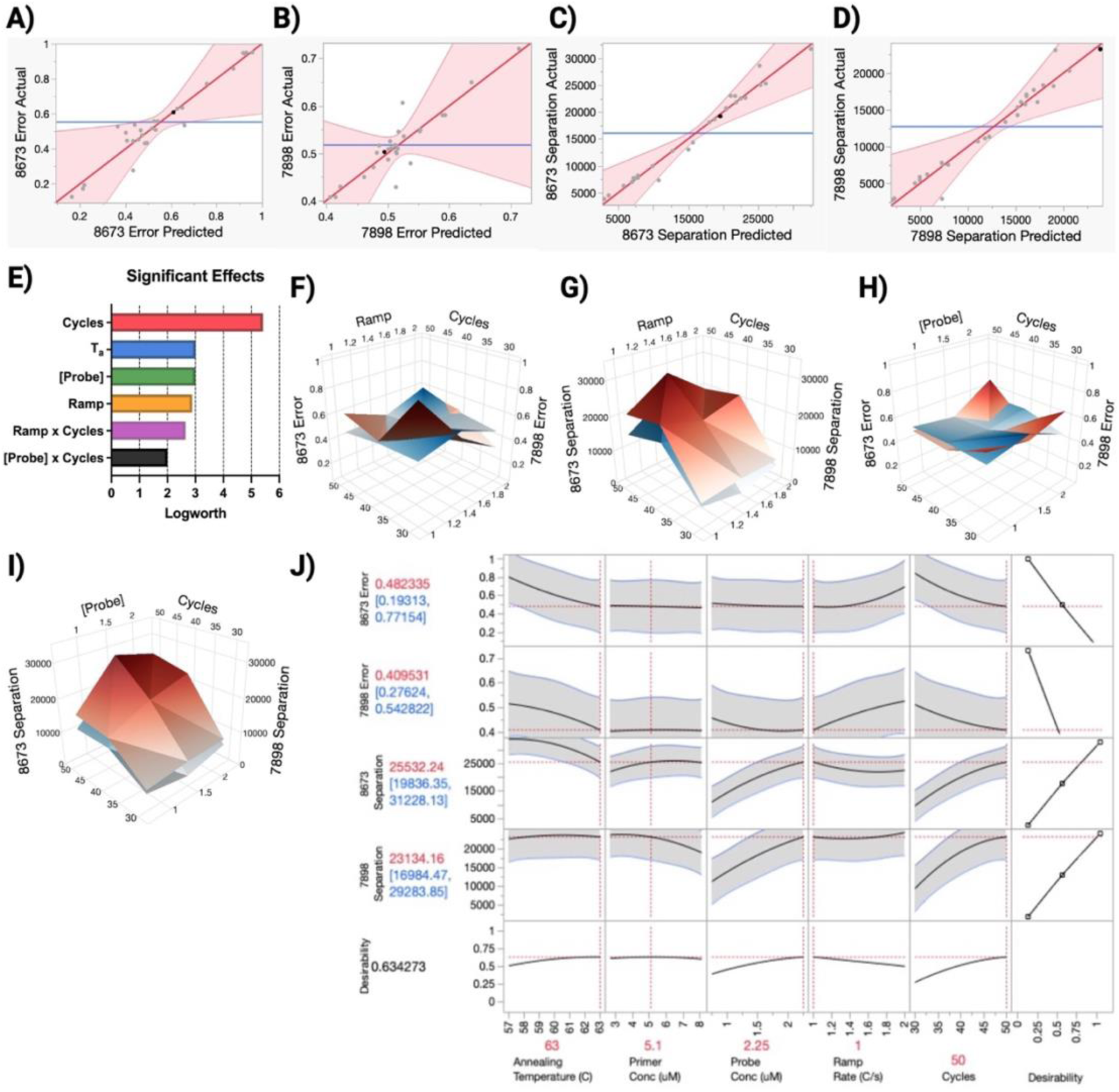
Central composite design modeling. Actual *vs.* predicted model fit to the relative error of the assay for **(A)** transgene (8673) and **(B)** genome (7898) barcodes. Actual *vs.* predicted model fit to the separation of mean fluorescence intensity between positive and negative droplet populations of the assay for **(C)** transgene and **(D)** genome barcodes, respectively. **(E)** Plots of the log-transformation of the effect significance of design factors (T_a_, annealing temperature; [Probe], probe concentration; Ramp, ramp rate). Response surface graphs of **(F)** relative error and **(G)** droplet separation as a function of the ramp rate x cycles; and **(H)** relative error and **(I)** droplet separation as a function of probe concentration x cycles (red, 8673; blue, 7898). **(J)** Plots of the maximized desirability function with respect to the 5 design factors (annealing temperature, primer concentration, probe concentration, ramp rate, and cycles) and 4 optimization goals (minimize relative error and maximize separation for the transgene and genome barcodes)

To determine factor settings that yield a balance of optimal relative error and droplet separation for both the 8673 and 7898 barcodes, we set a desirability function to minimize the relative errors and maximize the droplet separation for the 8673 and 7898 barcodes. A maximum desirability of 0.78 was achieved, corresponding to predicted relative errors of 48% and 41% and droplet separations of 25,737 and 22,889 a.u. for barcodes 8673 and 7898, respectively. The factor settings predicted to yield these values were a T_a_ = 63°C, a primer concentration = 5.1 μM, a probe concentration = 2.25 μM, a ramp rate of 1°C/s, and 50 PCR cycles (**Fig 2J**).

### The optimized ddPCR assay exhibits high precision and dynamic range after correction by linear regression

Having characterized the effects of the PCR settings on assay performance, we next proceeded to assess the performance of the assay using the settings predicted to optimize the desirability function (herein, optimized ddPCR). To do so, we prepared eight dilutions of the reference plasmids in 5-fold intervals, ranging from a 6.44×10^5^ to 8 copies/well. We then measured the concentration of these dilutions using optimized ddPCR and plotted against their predicted concentrations. In the dynamic range of the assay, we observed minimum relative errors of 51% and 43%, closely mirroring the 48% and 41% relative errors predicted by the least-squares models (**Fig 3A and 3B**). This error was observed with high precision, indicating that it is systematic error in the assay and, thus, we next sought to correct for this using a calibration curve.

**Fig 3.**
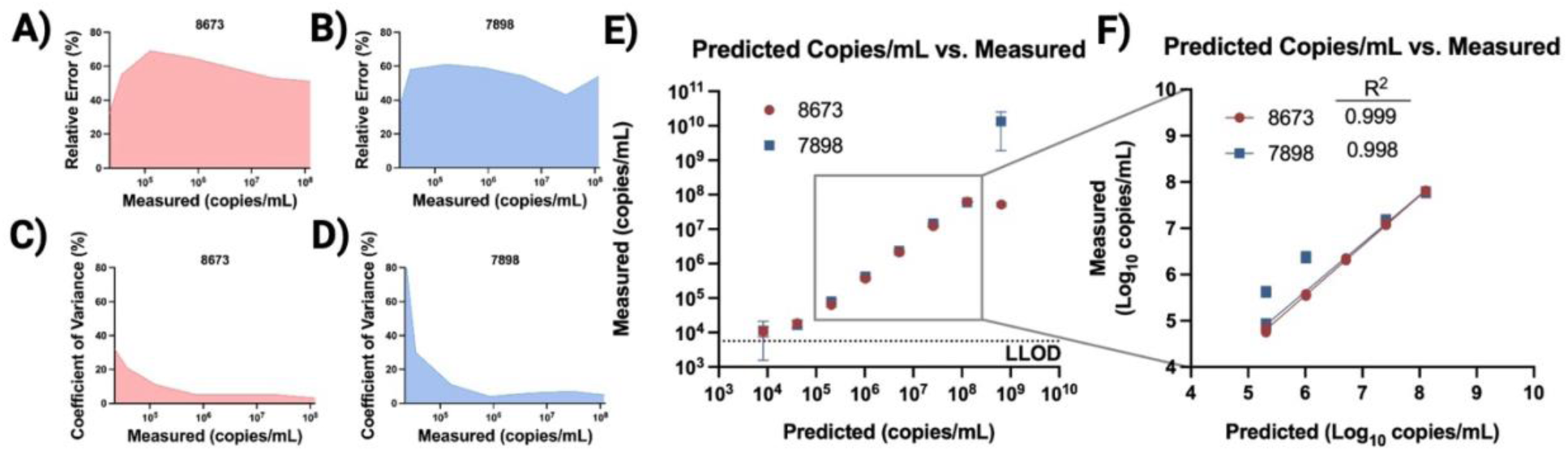
ddPCR assay performance characterization. The relative error of the ddPCR measured concentrations, as compared to the predicted concentrations, for the **(A)** transgene and **(B)** genome barcodes. The coefficient of variance as a function of ddPCR measured **(C)** transgene and **(D)** genome barcode concentrations. **(E)** Concentrations of transgene (8673) and genome (7898) barcode DNA quantified by Qubit (predicted) compared to concentrations quantified by ddPCR (measured) with the lower limit of detection (LLOD) denoted. **(F)** Linear regression analysis of five of the concentrations in **(E)** (8673 R^2^ = 0.999, slope = 1.08, 95% CI [1.06, 1.09], p < 0.0001, n = 27; 7898 R^2^ = 0.998, slope = 1.04, 95% CI [1.01, 1.07], p < 0.0001, n = 30). Points represent mean ± SD.

We found a linear relationship between measured and predicted values at concentrations between 63 – 62,620 and 80 – 59,469 measured copies/well, with a precision (defined as the coefficient of variance, CV) of 3-5% and 4-7% (**Fig 3C and 3D**), and a lower limit of detection (LLOD) of 5.66 and 6.66 copies/well, for 8673 and 7898 respectively (**Fig 3E**). These values correspond to a dynamic range of 6.4×10^4^ – 6.3×10^7^ and 8.0×10^4^ – 6.0×10^7^ measured copies/mL, with LLOD of 5,661 and 3,623 copies/mL, for 8673 and 7898, respectively **(Table 1**). Performing linear regression on the data in the dynamic range of the assay revealed strong fits, with R^2^ = 0.999 and 0.998 for 8673 and 7898, respectively (**Fig 3F**). With this strong fit, the following linear equations can be used to correct for the inherent error of the assay in the dynamic range of measured template concentrations, where C_a_ is the true concentration and C_m_ is the measured concentration (**equations 1 and 2**).

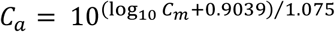

Equation 1. 8673 error correction

**Table 1.**
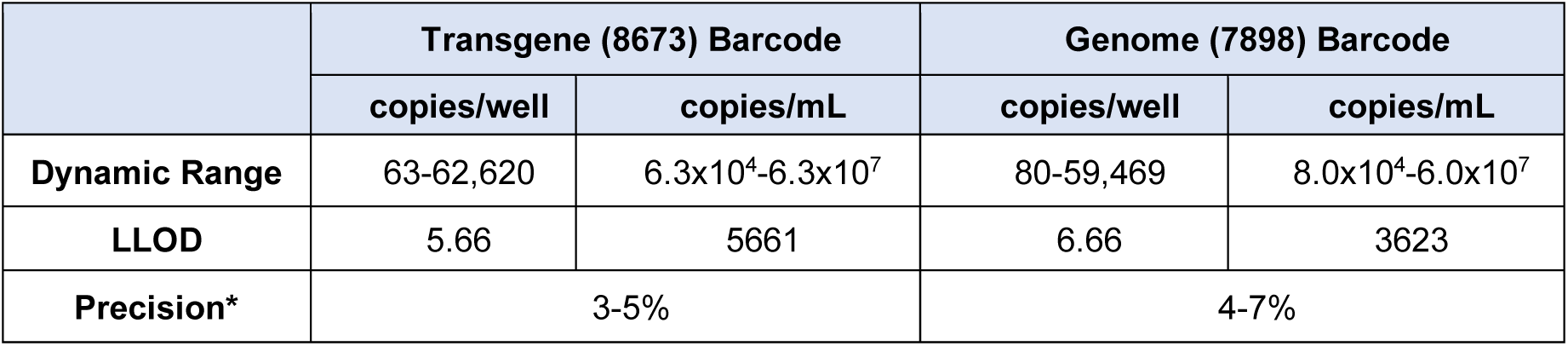
ddPCR assay performance metricst. *in the dynamic range

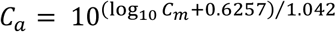

Equation 2. 7898 error correction

### Optimized ddPCR is a more accurate measure of phage vector titer and purity than biological activity assays

Currently, the de facto gold standard approach to assess the biological activity of a phage vector is via measuring the number of bacteria transduced or number of vector particles that can form plaques on host cells, which may not accurately reflect the true number of transgene-packed phage vector capsids present. To understand how the transgene:vector genome copies/mL titers compare to biological activity, we generated T7-based phage vectors using established methods [19,21,22]. Briefly, the 8673 barcode was inserted into a plasmid containing a T7 packing signal and a kanamycin resistance cassette, whereas the 7898 barcode was inserted in the T7 genome in place of gene 17, which encodes the T7 tail fiber. Removal of this gene prevents any vector capsids packed with the genome from establishing a productive infection as tail-fiber deficient T7 particles lack the host-receptor binding domain [23]. While tail-fiber competent vectors particles packed with the barcoded genomic DNA can still cause lysis if delivered to a target cell, the progeny from this infection are tail fiber deficient, and thus unable to infect subsequent cells. *E. coli* containing the barcoded transgene plasmid and a complementary plasmid expressing the T7 tail fiber were then infected by the barcoded T7 helper phage to generate the vectors.

Using the ddPCR assay optimized above, we first quantified the number of transgene and vector genome DNA packed phages, with the ratio of these representing the purity of the phage vector batch. The conventional process of generating T7 phage vectors involves infecting bacteria possessing transgenic DNA of interest with a helper phage, which contains only genome-packed capsids; the resulting T7 phage vector prep contains a mixture of both genome and transgene packed phage particles. When samples of either T7 helper phage or T7 phage vectors were analyzed, transgene-packed capsids were only detected in the vector sample, confirming high assay specificity (**Fig 4A**). Next, to test the extent to which free DNA contributes to the ddPCR signal, we spiked a helper phage sample with free transgene plasmid DNA. After DNA purification, including a DNase treatment step to remove non-encapsidated free DNA, we observed a reduction in transgene copies by >3 orders of magnitude, indicating that free DNA contributes to <0.1% of the detected DNA copies for measurements made following our preparation steps (**Fig 4B**). This confirms the ability of our assay and workflow to measure only encapsidated phage vector DNA.

**Fig 4.**
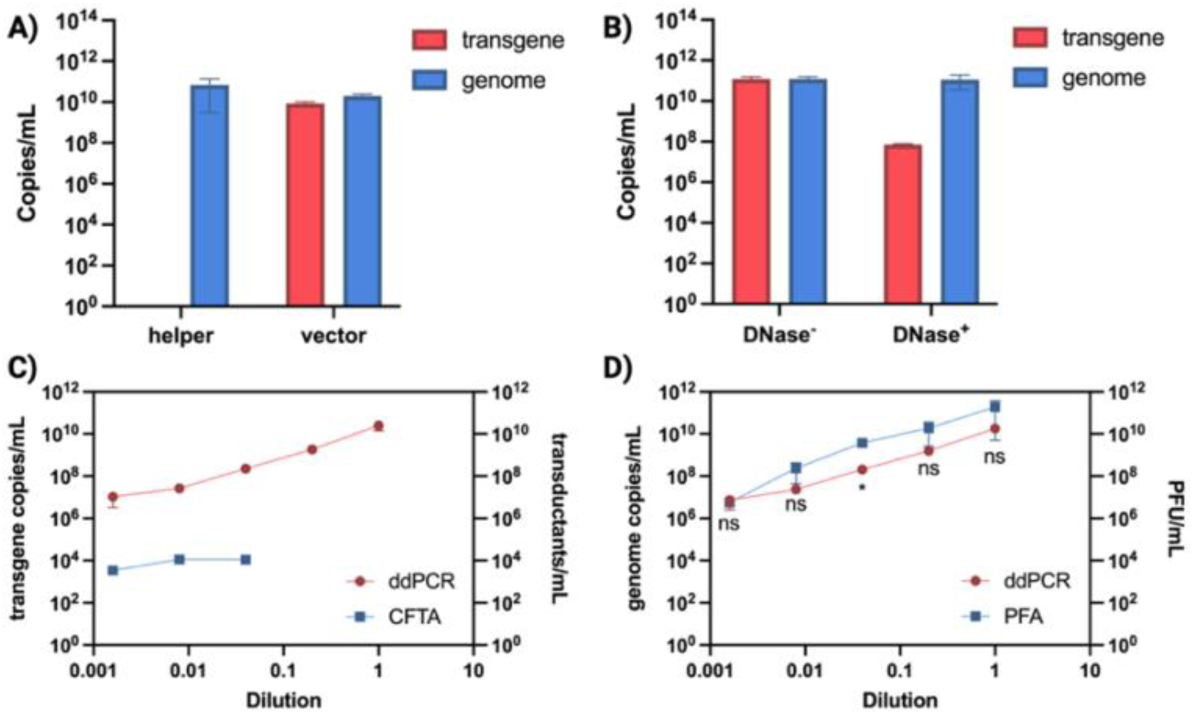
Characterization of phage vectors by ddPCR and biological activity assays. **(A)** Concentrations of transgene and genome copies in DNA purified from bacteriophage lysates determined by ddPCR (helper, T7 helper phage particles packed exclusively with barcoded genomic DNA; vector, T7 vector particles, packed with either barcoded transgenic DNA or genomic DNA. **(B)** ddPCR quantified transgene and genome copies with and without DNase treatment to digest non-encapsidated DNA during DNA purification of a helper phage DNA spiked with free plasmid DNA. **(C)** Comparison of transgene-packed vector particle quantification by colony formation transduction assay (CFTA) and ddPCR. **(D)** Comparison of genome-packed vector particle quantification by plaque formation assay (PFA) and ddPCR. Differences between ddPCR genome copies and plaque assay titers (PFA) across dilutions were analyzed using a two-way repeated-measures ANOVA (Šídák’s post-hoc comparison test; *p < 0.05; ns, not significant; n = 3)

We next assessed the vectors via biological activity assays, including colony formation transduction assays (CFTA), and by a trans-complementation plaque formation assay (PFA) using an *E. coli* strain with trans-complementary expression of the T7 tail fiber that allowed for plaque visualization. For both, we tested vectors across 5 different dilutions at 5-fold intervals. The CFTA showed colony formation for only the 3 most dilute vector samples and, notably, the colony counts at these 3 dilutions were nonlinear. This was strongly influenced by the toxicity of vector genome-packed particles, as vector genome-packed capsid titer correlated (R^2^ = 0.527, p = <0.0001) to target cell killing (**Supplementary Fig 1**). In comparison, our optimized ddPCR assay measured >3-orders of magnitude more transgene-packed vectors/mL than transductants at each vector concentration and was able to quantify the 2 vector samples that showed no transduction in the CFTA (**Fig 4C**). In contrast to the CFTA, there was no significant difference observed between the PFA and the ddPCR assay in 4 out of 5 of the vector dilutions (**Fig 4D**).

### ddPCR enables precise titering and purity assessment of temperate phage-derived vectors

To characterize the ability of the optimized ddPCR to accurately titer phage vectors derived from temperate—as opposed to lytic (e.g. T7)—phages, we generated a P1 phage vector production bacterial strain by first transforming an *E. coli* strain lysogenized by phage P1 with a transgene plasmid encoding i) the P1 packing signal (pacAB) and ii) a kanamycin resistance cassette (pacAB^+^). We then produced a variant of this strain that lacks pacAB in the P1 episome by replacing this locus with a chloramphenicol resistance cassette (pacAB^-^). Upon making P1 vectors from each of these strains, ddPCR titration revealed that vectors derived from the pacAB^+^ cell line contain a 1:1.3±0.1 ratio of vector genome- to transgene-packed capsids, whereas the pacAB^-^cell line produced vector preparations containing 166-fold more transgene-packed capsids than those packed with vector genome (**Fig 5A**). The precision afforded by the ddCPR assay also revealed that titer variability was dramatically lower between production runs when pacAB was knocked out in the P1 episome, with CVs of >100% for pacAB^+^ vectors vs. 15% for pacAB^-^ vectors (**Fig5B**).

**Fig 5.**
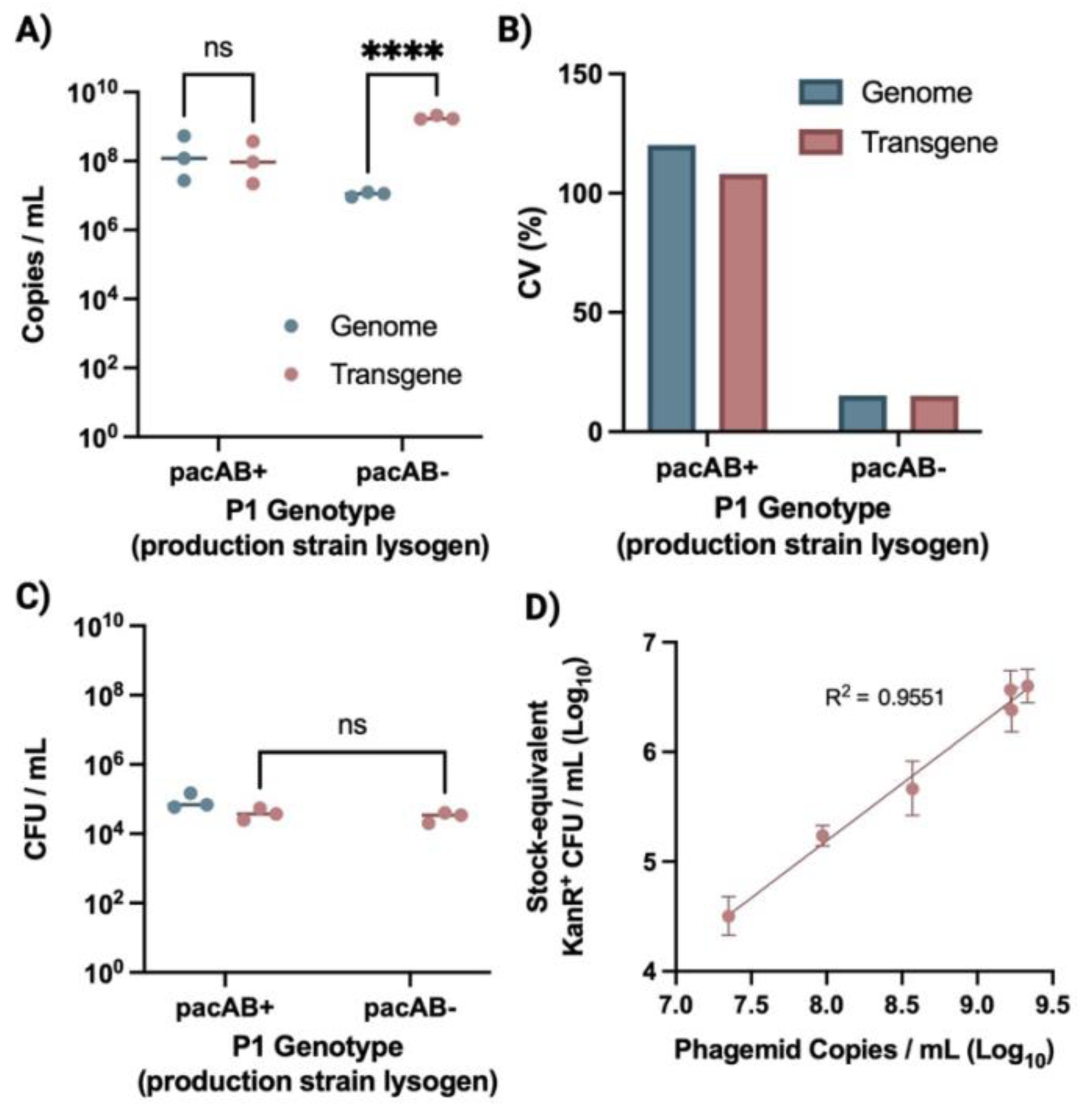
ddPCR-quantified packing composition, purity, and corresponding functional effects of temperate phage-derived vectors. **(A)** Vector genome- and transgene-packed capsid titers of P1 phage vectors produced from P1 production cell lines harboring either a packaging signal–competent episome (pacAB+) or a packaging signal-deficient episome (pacAB−). **(B)** Run-to-run variability for ddPCR titers expressed as coefficient of variation (CV) for genome- and phagemid-packed capsids in pacAB+ versus pacAB− vector preparations. **(C)** Transduction rates for pacAB+ versus pacAB− vector preparations when co-incubated with target cells at a multiplicity of infection (MOI) of 0.2 vectors per target cell, analyzed using a two-way repeated-measures ANOVA (Šídák’s post-hoc comparison test; ****p < 0.0001; ns, not significant; n = 3). **(D)** Relationship between ddPCR-measured transgene-packed capsids/mL and the number of cells transduced by these vectors, normalized to the dilution ratio used to achieve the MOI used in the transduction reaction (stock-equivalent KanR⁺ CFU/mL). The association between variables was evaluated using simple linear regression (R^2^ = 0.955, slope = 1.04, 95% CI [0.92, 1.16], p < 0.0001, n = 18). Points represent mean ± SD.

While these titer data equate to a purity of 99.4% transgene-packed capsids in vector lots derived from the pacAB^-^ cell line, there was still an average of 1.1 x 10^7^ vector genome-packed capsids/mL per batch. Despite the presence of these vector genome-packed capsids, no cells transduced by them were detected when a CFTA was performed at an MOI of 0.1 vectors per target cell (**Fig 5C**). However, when this MOI was increased to 0.2, low rates of transduction by genome-packed capsids were observed (**Supplementary Fig 2**), demonstrating that quantification by this optimized ddPCR assay provides critical insight into the composition and biological activity of vector batches. As expected, normalization of MOI to ddPCR-derived transgene-packed capsid titers results in transgene transduction rates that do not differ between pacAB^+^- and pacAB^-^-derived vectors (**Fig 5C**). Likewise, when transgene-transduced colony counts were normalized to the respective dilution used to achieve a constant MOI for each vector batch and plotted against their ddPCR-derived transgene-packed capsid titers, strong correlation was observed (R^2^ = 0.955) (**Fig 5D**), indicating the accuracy of the ddPCR assay in titering phage vectors derived from temperate phages, in addition to lytic derivatives.

### Optimized ddPCR is critical for producing high titer vectors and ensuring accurate and reproducible transduction

The titer of phage vector preparations depends, critically, on production reaction conditions, including the abundance of vector production bacteria, the amount of input helper phages as measured by multiplicity of infection (MOI), the medium, and other environmental conditions. Thus, to maximize the titer and efficacy of phage vector production, it is essential that we can accurately measure the titers and purity of different phage vector preps, as measured by transgene- and vector genome-packed vector particles, as opposed to biological activity assays, which can mis-represent titers (**Fig 4C**). Here, we used a full factorial experimental design to optimize transgene-packed vector titers by varying the cell density of the vector production strain culture, the titer of the helper phage inoculum, and the vector production incubation time after inoculation (**Supplementary Tables 6 and 7**). We then quantified number of both transgene- and genome-packed vector particles in each preparation. We found this spanned an 85-fold range, with a maximal transgene titer of 1.73×10^10^ copies/mL achieved (**Fig 6A & 6B**). By comparing the transgene- to genome-packed capsid titers, we found purity ratios (i.e. transgene:genome copies) varied from ∼4:1 to 1:1 (**Fig 6C**) and that a higher purity ratio is inversely correlated (r = −0.80, p = 0.009) to overall titer (**Supplementary Fig 3**).

**Fig 6.**
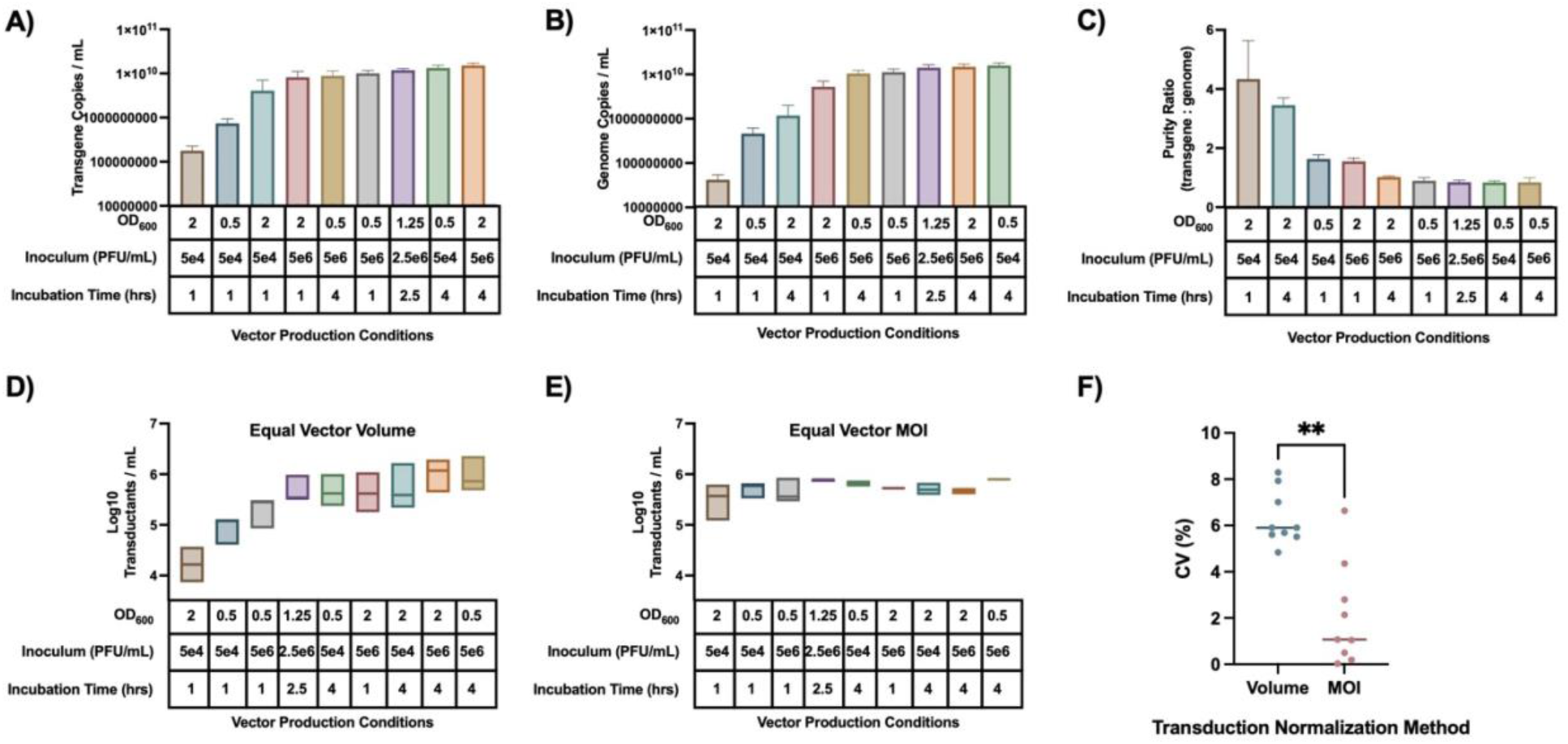
ddPCR titration facilitates vector production optimization and transduction reproducibility. **(A)** ddPCR-based transgene-packed vector particle titers as a function of vector production conditions. **(B)** ddPCR-based genome-packed vector particle titers as a function of vector production conditions. **(C)** Vector purity ratios (transgene-packed particles: genome-packed particles) as a function of production conditions. **(D)** Target bacteria that received cargo DNA from phage vectors (transductants)/mL as a function of vector production conditions. An equal volume of vectors was used for each transduction reaction. **(E)** Same as **D**, except vector volume was varied to achieve a constant MOI of 0.2 (1 vector per 5 target cells) for each transduction reaction. **(F)** Inter-replicate coefficients of variation (CV) calculated from log_10_-transformed transductant concentrations (**D** & **E**) for each vector production condition, comparing equal-volume versus equal-MOI transduction reaction normalization methods (two-tailed Mann–Whitney test; **p < 0.01; n = 9).

To demonstrate how titer data from the optimized ddPCR assay improves the accuracy and reproducibility of vector performance, we next performed transduction assays using either an equal volume of each vector preparation or, using the transgene- and genome-packed particle titer data, an equal MOI (dosing 1 transgene-packed particle for every 5 target cells). We observed that, without titer-informed MOI normalization, the number of cells that received DNA from the vectors (transductants) varied by more than 60-fold, ranging from 2×10^4^ – 1.2×10^6^ cells/mL (**Fig 6D**). In contrast, when titer data was used to normalize the transduction reaction to a constant MOI, transductants varied by less than 3-fold between vector preparations (**Fig 6E**) and the Log_10_-adjusted CV between replicates was also more than 300% lower when titering data was used to normalize the MOI (**Fig 5F**). Together, these data demonstrate that titering of distinct phage vector preparations using the optimized ddPCR assay is essential in determining optimized production conditions and facilitates more accurate and reproducible transduction, even across vector batches with different production conditions.

## Discussion

Similar to viral vectors used in human gene therapy, phage vectors represent a powerful new tool capable of genetically perturbing specific bacterial targets in complex microbial communities with precision, offering the potential to facilitate entirely new therapeutic paradigms, as well as causal gene-function perturbation studies within complex native communities *in situ*. To further their development, methods that can accurately characterize phage vectors are essential. Indeed, clinical use of viral vectors for human gene therapy requires extensive characterization of the viral vector preparations to ensure safety, potency, purity and stability. At the core of each of these categories is accurate and reliable quantitative measurements of the number of vector capsids packed with the DNA of interest, as well as the abundance of any contaminating proteins and DNA. Such information is critical to ensure reproducible efficacy, safety and limited immunogenicity, to correctly define the dose and/or dosing regimen, and to limit batch-to-batch variations in manufacturing as well as loss of activity during storage. The same characterizations are almost certainly required by regulatory authorities to advance phage vectors into human studies.

qPCR and ddPCR are commonly used to quantify copies of vector-packed DNA when characterizing viral vectors used in human gene therapy. While qPCR has traditionally been the more prevalent approach—as real-time thermal cyclers are more commonly available and less costly and time consuming to purchase, run, and maintain than ddPCR droplet generators and readers—ddPCR offers a number of advantages over qPCR. These include greater sensitivity of detection while simultaneously having less sensitivity to impurities that may disrupt PCR efficiency, and elimination of sources of error related to standard curves, which are not needed for absolute quantitation [24]. While ddPCR assays have been previously used to quantify naturally occurring lytic phages used as targeted antibiotics [25–27], there exist no established ddPCR or qPCR assays for quantification of phage vectors and assessment of their purity. We thus sought to advance here a ddPCR-based assay to address this gap.

Given the large number of input variables, we elected to utilize CCD to optimize a ddPCR assay for phage vector quantitation. Such response surface methodology has previously proven effective in optimizing a ddPCR assay for genetically modified crop certified reference materials [28]. We applied CCD to understand how the assay sensitivity is impacted by changes in PCR annealing temperature, primer and probe concentration, cycle ramp rate, and the number of PCR cycles, and to determine theoretically optimal settings for these factors. We found that, although less sensitive to changes in PCR conditions than droplet separation, PCR cycle number and annealing temperature strongly influence assay accuracy. Using the CCD-optimized settings, our ddPCR assay offers a dynamic range of nearly 3 orders of magnitude for both the 8673 and 7898 barcodes (987-fold and 745-fold, respectively). Within the dynamic range, the high degree of correlation between measured and known barcode concentrations allows calibration using the corresponding equations of the regression lines (**equations 1 and 2**) to correct for the systematic error of the assay, allowing for highly accurate titer quantitation of diverse types of phage vectors.

Phage vectors can be classified by both the underlying phage type (lytic or temperate) and their replication competency. While vector production systems have been described for temperate phages that yield vectors packed primarily with the transgenic DNA of interest [29], many phage vectors to date, including all lytic phage-based systems, comprise of a mixed population of both transgene and genome-packed vectors [10,21,22,30,31]. The data presented here demonstrate the utility of the optimized ddPCR assay in characterizing non-replicative vectors derived from both lytic and temperate phages. Following an established design and production method for replication-impaired (and thus at reduced risk of horizontal gene transfer) vectors derived from the lytic phage T7 [21,30], we show that a significant fraction of these vector batches consists of vector genome-packed capsids, which can induce considerable lysis of target bacteria. This underscores the importance of an assay that can accurately enumerate both populations, which we achieved here with high accuracy and sensitivity through barcodes embedded in the transgene-encoding phagemid and the vector genome.

We note important differences in titer estimates derived from conventional biological assays such as CFTAs and PFAs *vs.* the ddPCR assay. For instance, ddPCR quantified transgene-packed capsids at all 5 T7-based vector concentrations tested, whereas the CFTA only yielded transductants at the 3 most dilute concentrations, which was strongly influenced by target cell-directed toxicity associated with a higher number of vector genome-packed particles. Importantly, the number of transductants measured was far lower than the actual number of transgene-packed T7 vectors determined by ddPCR at each of these concentrations. Conversely, the genome-packed titers revealed by the ddPCR assay closely mirrored those given by the PFA—with any variation between the two likely due to inefficiencies in the DNA purification process and errors inherent to both the PFA and ddPCR assay—indicating that our optimized ddPCR assay is accurate with purified vector DNA in addition to reference plasmids.

Similarly, optimized ddPCR titration of non-replicative vectors derived from the temperate phage P1 showed comparable accuracy, as shown by the high degree of correlation (R^2^ = 0.955) between the number of cells transduced by P1 vectors and the volume of the vectors dosed, as determined by ddPCR. Comparison of P1 vectors derived from production cell lines with either packing signal competent or deficient vector genomes also revealed that packing signal knock out in the vector genome dramatically improves vector purity, reducing contamination by vector genome-packed capsids from 57% to 0.6%. When assessing the biological activity of these vectors, transduction by the low-abundance vector genome-packed capsids were only detected above a threshold MOI, highlighting the utility of this assay in both characterizing lytic and temperate phage-derived vectors, as well as understanding their phenotypic effects and biological activity.

With this assay, there are a number of experimental procedures critical to ensuring accurate quantitation, including DNase treatment during the DNA purification step. This reduces free, non-encapsidated DNA by more than 3 orders of magnitude, making the latter insignificant in calculating the final phage titers. In addition, to maximize assay precision and minimize the LLOD, we found it critical to minimize freeze-thaw cycles of all reagents, and to prepare the assay under a local fume extractor to reduce risk of contamination by aerosols.

The insights described here underscore why precise quantification of both transgene and vector genome-packed capsids in phage vector preparations is essential to a complete understanding of the vector system. In concert with data from biological activity assays, ddPCR readouts can help determine vector dosing, guide vector production, and ensure highly reproducible use both in preclinical and clinical studies. By carefully defining the error and precision of this assay, it can be used to quantify both the transgene- and genome-packed titers of phage vector samples with higher throughput and greater accuracy that biological activity assays alone. Furthermore, this work can be readily applied to a wide range of both lytic- and temperate-based phage vector systems of interest through the modular, highly orthogonal barcodes developed here. As such, this quantification method is well suited to support the development of next-generation phage vector systems, including those with advanced methods of vector production [32], allowing for more precise and reproducible transduction of a wide range of bacterial targets. With phage vectors poised to become versatile tools for microbiome engineering, we believe the approach reported here will have significant value as a key assessment standard.

## Statements and Declarations

### Author Contributions

Conceptualization, P.J.V. and S.K.L.; Methodology, P.J.V.; Investigation, P.J.V., R.P., and J.L.; Formal Analysis, P.J.V.; Writing – Original Draft, P.J.V.; Writing – Review & Editing, P.J.V. and S.K.L.; Funding Acquisition, S.K.L.

## Acknowledgements

Figures were created with BioRender.com and GraphPad Prism.

## Data Availability

The datasets generated and/or analyzed during the current study are available within the article and its Supplementary Information files, and at https://doi.org/10.5281/zenodo.17095292. Additional files may be available from the corresponding author upon reasonable request.

## Clinical Trial Number

not applicable.

