## Supplementary Data for "A Droplet Digital PCR Assay for Quantification of Bacteriophage Viral Vector Titer and Purity"

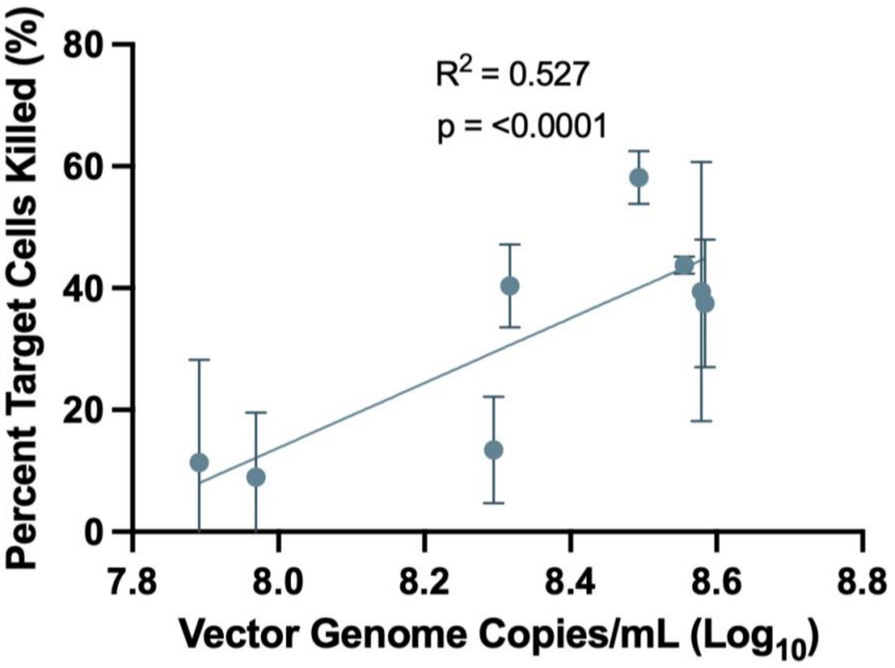


**Supplementary Figure 1. Target cell killing correlates to vector genome-packed capsid titers.** The percent of total target cells killed after coincubation with T7 phage vectors plotted against the $Log_{10}$ of the vector genome-packed capsid titer (copies/mL) of the T7 phage vector batch ($R^2$,p = correlation coefficient, significance value. n = 3).

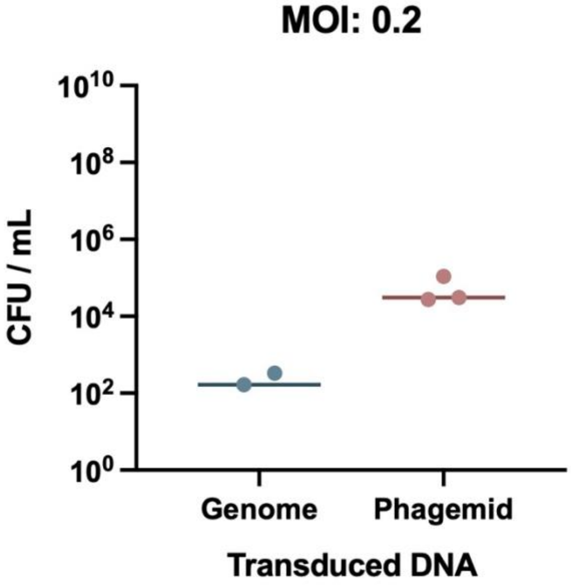


**Supplementary Figure 2. pacAB$^{-}$-derived P1 vector transduction at an MOI of 0.2.** Cells transduced by either vector genome or phagemid DNA via a colony formation transduction assay (CFTA) performed at a multiplicity of infection (MOI) of 0.2 vectors per target cell for P1 vectors derived from a production cells line containing a P1 episome encoding a chloramphenicol resistance cassette in place of the pacAB locus (n = 3).

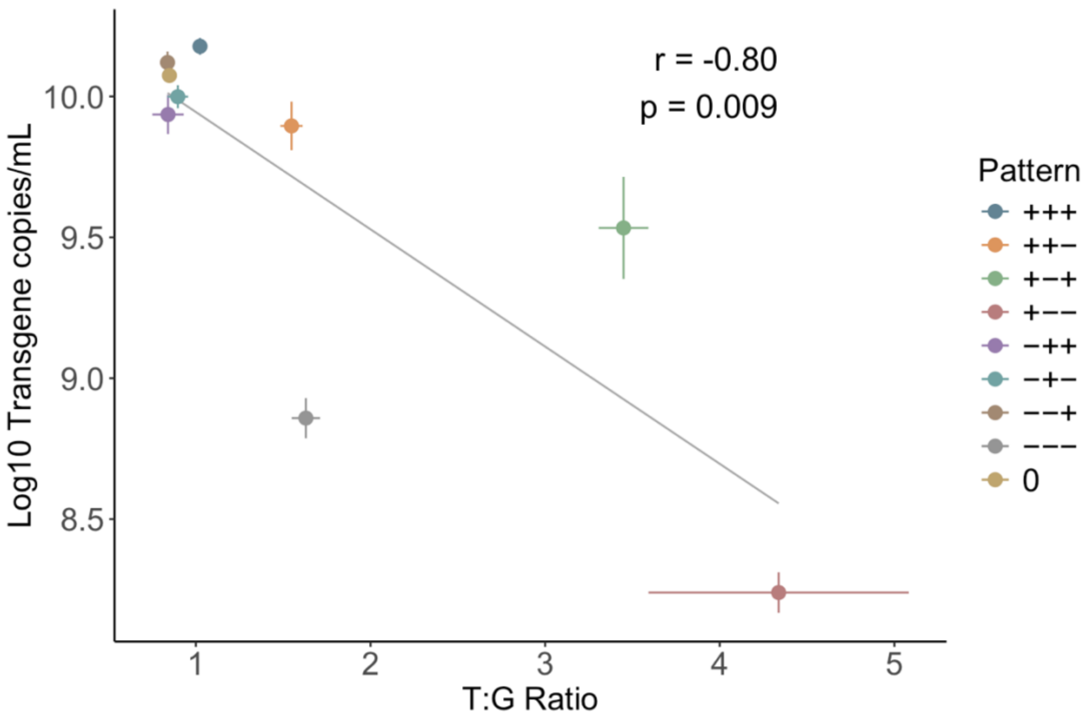


**Supplementary Figure 3. Purity ratio is inversely correlated with transgene titer.** Correlation plots of the purity ratio (transgene-packed vector particles / mL to genome-packed vector particles / mL; T:G Ratio) vs. $Log_{10}$ transgene copies / mL of phage vector preparations manufactured under varied production conditions. The association between variables was evaluated using Pearson's correlation coefficient (r = −0.80, 95% CI [-0.96, -0.30], two-tailed p = 0.009, n = 9 pattern means). Points represent the mean ± SE of three independent preparations per pattern. See **Supplementary Table 5** for production condition details corresponding to each pattern group.

| Barcode | Sequence |
| --- | --- |
| 8673 | ACTTCCATCGCTTCTCTTGGGACGTTTCTTAACTGCTGCAAGACTACTTGAACATTTCGACTTATAATCTATCCGATACGCTAGTTGGGCTGCTCAGACTACCAGTCC<br>CGTTCATTGCCATCCCCGTCCAATTTAAGAACGTGTATGATATGCGAAGCACCTGAAATACCTAGTACCTAATACTCCTACAGCCCCTTTATTGTCTTACCGTATTCC<br>GACCTCTACACCGAGGGGTACGTTAATTCGCCAAAACGATCTTGAGTGGCAATATGCCAACTTGAATACCGTGCCACCGTTACA |
| 7898 | GGCGGGGTTGAACGGTCAGGTATCACTATAATCAAAACCCCTCGATATATGCATTAATCTCGCCGGTCCGATGAACTGACGGTCATGTAGCTAGTATGAGCCGCG<br>AAGAGGGACGGAGACATGCCAAACGCGAACAAGGTCATCCCTATGCCGAACAATATGTCGCCCTGGCTTTATAGCAGATTACGGTCAGCGCCGCTATATACATA<br>AACTGTGACTAACATATTATCCCTGTGTATAACACCAGTTCGATCCGAAACCCAAATGTATCAGAGTCACCGTTAATCCACTAAAGGTCTG |

**Supplementary Table 1. Barcode sequences.** Barcode 8673 was inserted into the DNA containing the transgene cassette. Barcode 7898 was inserted into the T7 genomic DNA.

| Barcode | Secondary Structure | | Top BLAST Result | | | Orthogonality† |
|---|---|---|---|---|---|---|
| | Max MFE* | Avg MFE* | Alignment Length | Percent Identity | Query Coverage | |
| 8673 | -10.9 | -5.09 | 28 | 96.4 | 9 | 250 |
| 7898 | -11.8 | -5.663 | 24 | 100 | 8 | 250 |

**Supplementary Table 2. Barcode metrics.** *MFE values are the greatest of those calculated for a 50 bp sliding window over the full sequence. †Orthogonality as defined by Hamming distance.

| Factor | Low | Central | High |
|---|---|---|---|
| $T_a$ (°C) | 57 | 60 | 63 |
| [Primer] (µM) | 2.7 | 5.4 | 8.1 |
| [Probe] (µM) | 0.75 | 1.5 | 2.25 |
| Ramp Rate (°C/s) | 1 | 1.5 | 2 |
| Cycles | 30 | 40 | 50 |

**Supplementary Table 3. ddPCR central composite design factors.** Factors and corresponding low (-), central (0), and high (+) settings used in the central composite design (CCD) used to optimize the performance of the ddPCR assay. Factors include annealing temperature ($T_a$), primer concentration ([Primer]), probe concentration ([Probe]), PCR thermal cycling ramp rate, and number of amplification cycles. Optimization targeted minimal relative error and maximal separation of positive and negative droplet populations for both the transgene- (8673) and genome-encoded (7898) barcodes.

| Pattern | Annealing Temperature (C) | Primer concentration (nM) | Probe concentration (nM) | Ramp Rate (C/s) |
|---|---|---|---|---|
| 00a00 | 60 | 5400 | 750 | 1.5 |
| 0000a | 60 | 5400 | 1500 | 1.5 |
| 000a0 | 60 | 5400 | 1500 | 1 |
| 0 | 60 | 5400 | 1500 | 1.5 |
| +−+++ | 63 | 2700 | 2250 | 2 |
| +−−+− | 63 | 2700 | 750 | 2 |
| −−−−− | 57 | 2700 | 750 | 1 |
| −++++ | 57 | 8100 | 2250 | 2 |
| +−+−− | 63 | 2700 | 2250 | 1 |
| ++++− | 63 | 8100 | 2250 | 2 |
| 00A00 | 60 | 5400 | 2250 | 1.5 |
| 000A0 | 60 | 5400 | 1500 | 2 |
| +−−−+ | 63 | 2700 | 750 | 1 |
| ++−−− | 63 | 8100 | 750 | 1 |
| 0000A | 60 | 5400 | 1500 | 1.5 |
| −−−++ | 57 | 2700 | 750 | 2 |
| 0 | 60 | 5400 | 1500 | 1.5 |
| 0a000 | 60 | 2700 | 1500 | 1.5 |
| −++−− | 57 | 8100 | 2250 | 1 |
| a0000 | 57 | 5400 | 1500 | 1.5 |
| −−++− | 57 | 2700 | 2250 | 2 |
| −+−−+ | 57 | 8100 | 750 | 1 |
| 0A000 | 60 | 8100 | 1500 | 1.5 |
| −−+−+ | 57 | 2700 | 2250 | 1 |
| +++−+ | 63 | 8100 | 2250 | 1 |
| A0000 | 63 | 5400 | 1500 | 1.5 |
| −+−+− | 57 | 8100 | 750 | 2 |
| ++−++ | 63 | 8100 | 750 | 2 |

**Supplementary Table 4. ddPCR central composite design conditions.** Complete experimental conditions for the central composite design (CCD) used to optimize the ddPCR assay. Each row specifies individual runs with varying combinations of annealing temperature, primer concentration, probe concentration, and ramp rate.

| Pattern | Annealing Temperature (C) | Primer concentration (nM) | Probe concentration (nM) | Ramp Rate (C/s) |
|---|---|---|---|---|
| 00a00 | 0.491800131 | 0.549147334 | 14315.12 | 11344.37 |
| 0000a | 0.557776622 | 0.60634253 | 7276.65 | 2908.37 |
| 000a0 | 0.534136931 | 0.516468723 | 22487.25 | 17686.04 |
| 0 | 0.276726146 | 0.428413322 | 20492.72 | 17068.2 |
| +−+++ | 0.608292699 | 0.502621282 | 19208.4 | 23248.98 |
| +−−+− | 0.508798389 | 0.546252119 | 3815.26 | 2756.97 |
| −−−−− | 0.950143107 | 0.580407165 | 7655.85 | 5012.11 |
| −++++ | 0.857681377 | 0.720385693 | 25270.01 | 20296.25 |
| +−+−− | 0.946945538 | 0.486487262 | 6249.98 | 7563.8 |
| ++++− | 0.51485008 | 0.449695779 | 7696.09 | 7805.93 |
| 00A00 | 0.431575022 | 0.545275553 | 22963.79 | 18328.69 |
| 000A0 | 0.52760038 | 0.509710872 | 21746.24 | 16046.06 |
| +−−−+ | 0.556789518 | 0.508459742 | 7740.08 | 11109.63 |
| ++−−− | 0.948646262 | 0.580610766 | 4465.35 | 6212.24 |
| 0000A | 0.477446742 | 0.478665277 | 28593.92 | 23103.75 |
| −−−++ | 0.172783387 | 0.649692504 | 22642.26 | 15759.14 |
| 0 | 0.444070243 | 0.502589903 | 20715.08 | 16843.21 |
| 0a000 | 0.446685911 | 0.516221881 | 18051.65 | 15345.96 |
| −++−− | 0.941639061 | 0.499643626 | 9914.83 | 5452.66 |
| a0000 | 0.537151365 | 0.517224989 | 22987.15 | 14275.67 |
| −−++− | 0.193313123 | 0.407397891 | 6956.06 | 2961.1 |
| −+−−+ | 0.628754002 | 0.429551998 | 19150.97 | 7534.07 |
| 0A000 | 0.453283606 | 0.525159205 | 18623.99 | 13347.63 |
| −−+−+ | 0.771318935 | 0.510704065 | 31709.58 | 23171.13 |
| +++−+ | 0.505825127 | 0.40719496 | 25001.51 | 18207.61 |
| A0000 | 0.433778661 | 0.500899399 | 13005.04 | 16049.03 |
| −+−+− | 0.126162892 | 0.470395694 | 8160.58 | 5866.09 |
| ++−++ | 0.634819296 | 0.535697372 | 12730.81 | 10710.7 |

**Supplementary Table 5. ddPCR central composite design results.** Readouts for each set of factors for the central composite design (CCD) used to optimize the ddPCR assay. Each row to the design patterns detailed in Supplementary Table 4.

| Factor | Low | Central | High |
|---|---|---|---|
| Production Cell Density (OD600) | 0.5 | 1.25 | 2 |
| Helper Phage Inoculum (PFU/mL) | 5.00E+04 | 2.53E+06 | 5.00E+06 |
| Production Incubation Time (hrs) | 1 | 2.5 | 4 |

**Supplementary Table 6. Full factorial design of phage vector production optimization parameters.** Optimization parameters and their associated low (-), central (0), and high (+) levels used in the fractional factorial design for optimizing phage vector production conditions, including cell density, helper phage inoculum concentration, and incubation time.

| Pattern | Production Cell Density (OD600) | Helper Phage Inoculum (PFU/mL) | Production Incubation Time (hrs) |
|---|---|---|---|
| −−− | 0.5 | 5.00E+04 | 1 |
| −−+ | 0.5 | 5.00E+04 | 4 |
| −+− | 0.5 | 5.00E+06 | 1 |
| −++ | 0.5 | 5.00E+06 | 4 |
| +−− | 2 | 5.00E+04 | 1 |
| +−+ | 2 | 5.00E+04 | 4 |
| ++− | 2 | 5.00E+06 | 1 |
| +++ | 2 | 5.00E+06 | 4 |
| 0 | 1.25 | 2.53E+06 | 2.5 |

**Supplementary Table 7. Full factorial design phage vector production runs.** Detailed experimental conditions used for fractional factorial design runs, specifying the combinations of production cell density, helper phage inoculum, and incubation time applied to optimize phage vector production efficiency.

| Name | Sequence |
|---|---|
| PJV1063 | TTACTTACGACACTCCGAGACAGTCAGAGGGTATTTATTGAACTAGCATAAATAGGCGTATCACGAGGCAGAATTTCAGATAAAAAAAATCCTTAGCTTTCGCTAAGGAT<br>GATTTCTGGAATTCGCGGCCGCTTCTAGAGACTAGTGGAAGACATGGAGCGTAGGAAATAATACGACTCACTATAGGGAGAGGCGAAATAATCTTCTCCCTGTAGTC<br>TCTTAGATTTACTTTAAGGAGGTCAAATGATGGCTAACGTAATTAAAACCGTTTTGACTTACCAGTTAGATGGCTCCAATCGTGATTTTAATATCCCGTTTGAGTATCTA<br>GCCCGTAAGTTCGTAGTGGTAACTCTTATTGGTGTAGACCGAAAGGTCCTTACGATTAATACAGACTATCGCTTTGCTACACGTACTACTATCTCTCTGACAAAGGCTT<br>GGGGTCCAGCCGATGGCTACACGACCATCGAGTTACGTCGAGTAACCTCCACTACCGACCGATTGGTTGACTTTACGGATGGTTCCATCCTCCGCGCGTATGACCT<br>TAACGTCGCTCAGATTCAAACGATGCACGTAGCGGAAGAGGCCCGTGACCTCACTACGGATACTATCGGTGTCAATAACGATGGTCACTTGGATGCTCGTGGTCGTC<br>GAATTGTGAACCTAGCGAACGCCGTGGATGACCGCGATGCTGTTCCGTTTGGTCAACTAAAGACCATGAACCAGAACTCCTGGCAAGCACGTAATGAAGCCTTACAG<br>TTCCGTAATGAGGCTGAGACTTTCAGAAACCAAGCGGAGGGCTTTAAGAACGAGTCCAGTACCAACGCTACGAACACAAAGCAGTGGCGCGATGAGACAAAGGGTTT<br>CCGAGACGAAGCCAAGCGGTTCAAGAATACGGCTGGTCAATACGCTACATCTGCTGGGAACTCTGCTTCCGCTGCGCATCAATCTGAGGTAAACGCTGAGAACTCTG<br>CCACAGCATCCGCTAACTCTGCTCATTTGGCAGAACAGCAAGCAGACCGTGCGGAACGTGAGGCAGACAAGCTGGAAAATTACAATGGATTGGCTGGTGCAATTGAT<br>AAGGTAGATGGAACCAATGTGTACTGGAAAGGAAATATTCACGCTAACGGGCGCCTTTACATGACCACAAACGGTTTTGACTGTGGCCAGTATCAACAGTTCTTTGGT<br>GGTGTCACTAATCGTTACTCTGTCATGGAGTGGGGAGATGAGAACGGATGGCTGATGTATGTTCAACGTAGAGAGTGGACAACAGCGATAGGCGGTAACATCCAGT<br>TAGTAGTAAACGGACAGATCATCACCCAAGGTGGAGCCATGACCGGTCAGCTAAAATTGCAGAATGGGCATGTTCTTCAATTAGAGTCCGCATCCGACAAGGCGCAC<br>TATATTCTATCTAAAGATGGTAACAGGAATAACTGGTACATTGGTAGAGGGAGCGATAACAACAATGACTGTACCTTCCACTCCTATGTACATGGTACGACCTTAACACT<br>CAAGCAGGACTATGCAGTAGTTAACAAACACTTCCACGTAGGTCAGGCCGTTGTGGCCACTGATGGTAATATTCAAGGTACTAAGTGGGGAGGTAAATGGCTGGATG<br>CTTACCTACGTGACAGCTTCGTTGCGAAGTCCAAGGCGTGGACTCAGGTGTGGTCTGGTAGTGCTGGCGGTGGGGTAAGTGTGACTGTTTCCCAGGATCTCCGCT<br>TCCGCAATATCTGGATTAAGTGTGCCAACAACTCTTGGAACTTCTTCCGTACTGGCCCCGATGGAATCTACTTCATAGCCTCTGATGGTGGATGGTTACGATTCCAAA<br>TACACTCCAACGGTCTAGGATTCAAGAATATTGCAGACAGTCGTTCCGTACCTAATGCAATCATGGTGGAGAACGAGTAAAGGTGGCCAATTATTGAAGGCCTCCCTA<br>ACGGGGGGCCTTTTTTTGTTTCTGGTCTGCCCTCGCTTATGTCTTCTACTAGTAGCGGCCGCTGCTCCGGCAAAAAAGGGCAAGGTGTCACCACCCTGCCCTTTTT<br>CTTTAAAACCGAAAAGAATCGGTGTGAAAAGTCAGTATCCAGTCGTGTAGTTCTTATTACCT |
| PJV1161 | ATCGGTGTGAAAAGTCAGTATCCAGTCGTGTAGTTCTTATTACCTCGCGGAACCCCTATTTGTTTATTTTTCTAAATACATTCAAATATGTATCCGCTCATGAGACAATA<br>ACCCTGATAAATGCTTCAATAATATTGAAAAAGGAAGAGTATGAGTATTCAACATTTCCGTGTCGCCCTTATTCCCTTTTTTGCGGCATTTTGCCTTCCTGTTTTTGCTC<br>ACCCAGAAACGCTGGTGAAAGTAAAAGATGCTGAAGATCAGTTGGGTGCCCGAGTGGGTTACATCGAACTGGATCTCAACAGCGGTAAGATCCTTGAGAGTTTTCGC<br>CCCGAAGAACGTTTTCCAATGATGAGCACTTTTAAAGTTCTGCTATGTGGCGCGGTATTATCCCGTATTGACGCCGGGCAAGAGCAACTCGGTCGCCGCATACACTA<br>TTCTCAGAATGACTTGGTTGAGTACAGCCCAGTCACAGAAAAGCATCTTACGGATGGCATGACAGTAAGAGAATTATGCAGTGCTGCCATAACCATGAGTGATAACAC<br>TGCGGCCAACTTACTTCTGACAACGATCGGAGGACCGAAGGAGCTAACCGCTTTTTTGCACAACATGGGGGATCATGTAACTCGCCTTGATCGTTGGGAACCGGAG<br>CTGAATGAAGCCATACCAAACGACGAGCGTGACACCACGATGCCTGTAGCAATGGCAACAACGTTGCGCAAACTATTAACTGGCGAACTACTTACTCTAGCTTCCCGG<br>CAACAATTAATAGACTGGATGGAGGCGGATAAAGTTGCAGGACCACTTCTGCGCAGTGCCCTTCCGGCTGGCTGGTTTATTGCTGATAAATCCGGCGCTGGCGAAA<br>GAGGTTCTCGTGGCATAATCGCCGCGCTTGGTCCAGATGGTAAGCCCTCCCGTATCGTAGTTATCTACACGACGGGGAGTCAGGCAACTATGGATGAACGAAATAG<br>ACAGATCGCTGAGATAGGTGCCTCCCTGATTAAGCATTGGTAACTGTCAGACCAAGTTTACTCATATATACTTTAGATTGATTTAAAACTTCATTTTTAATTTAAAAGGAT<br>CTAGGTGAAGATCCTTTTTGATAATCTCATGACCAAAATCCCTTAACGTGAGTTTTCGTTCCACTGAGCGTCAGACCCCGTAGAAAAGATCAAAGGATCTTCATCACGG<br>CACTACACTCGTTGCTTTATCGGTATTGTTATTACAGA |
| PJV720 | ATCACGGCACTACACTCGTTGCTTTATCGGTATTGTTATTACAGATTTCCATAGGCTCCGCCCCCCTGACAAGCATCACGAAATCTGACGCTCAAATCAGTGGTGGCG<br>AAACCCGACAGGACTATAAAGATACCAGGCGTTTCCCCCTGGCGGCTCCCTCGTGCGCTCTCCTGTTCCTGCCTTTCGGTTTACCGGTGTCATTCCGCTGTTATGG<br>CCGCGTTTGTCTCATTCCACGCCTGACACTCAGTTCCGGGTAGGCAGTTCGCTCCAAGCTGGACTGTATGCACGAACCCCCCGTTCAGTCCGACCGCTGCGCCTTA<br>TCCGGTAACTATCGTCTTGAGTCCAACCCGGAAAGACATGCAAAAGCACCACTGGCAGCAGCCACTGGTAATTGATTTAGAGGAGTTAGTCTTGAAGTCATGCGCCG<br>GTTAAGGCTAAACTGAAAGGACAAGTTTTGGTGACTGCGCTCCTCCAAGCCAGTTACCTCGGTTCAAAGAGTTGGTAGCTCAGAGAACCTTCGAAAAACCGCCCTGC<br>AAGGCGGTTTTTTCGTTTTCAGAGCAAGAGATTACGCGCAGACCAAAACGATCTCAAGAAGATCCTTTGATCTTTTCTACGGGGTCTGACGCTTACTTACGACACTCC<br>GAGACAGTCAGAGGGTATTTATTGAACTA |
| PJV684 | ACCCACGACTATTGACTGCTCTGAGAAAGTTGATTGTTACGATTACATTGCTGGTGGTACAAGGAGGTCGAGGGTGAAGTACTTGCTGACTTCCTTGAGGAACACAT<br>GATGCGTCCTACGGTTGCTGCTACGCATATCATTGAGATGTCTGTGGGAGGAGTTGATGTGTACTCTGAGGACGATGAGGGTTACGGTACGTCTTTCATTGAGTGG<br>TGATTTATGCATTAGGACTGCATAGGGATGCACTATAGACCACGGATGGTCAGTTCTTTAAGTTACTGAAAAGACACGATAAATTAATACGACTCACTATAGGGAGAGG<br>AGGGACGAAAGGTTACTATATAGATACTGAATGAATACTTATAGAGTGCATAAAGTATGCATAATGGTGTACCTAGAGTGACCTCTAAGAATGGTGATTATATTGTATTA<br>GTATCACCTTAACTTAAGTTCCTATAGGGTCCTTTAAAATATACCATAAAAATCTGAGTGACTATCTCACAGTGTACGGACCTAAAGTTCCCCCATAGGGGGTACCTAAA<br>GCCCAGCCAATCACCTAAAGTCAACCTTCGGTTGACCTTGAGGGTTCCCTAAGGGTTGGGGATGACCCTTGGGTTTGTCTTTGGGTGTTACCTTGAGTGTCTCTCT<br>GTGTCCCTATCTGTTACAGTCTCCTAAAGTATCCTCCTAAAGTCACCTCCTAACGCTCGAGGCCCTATCTCCTTCGTCCACCAGCAATGTTACTTACGACACTCCGAGA<br>CAGTCAGAGGGTATTTATTGAACTA |
| PJV686 | TTACTTACGACACTCCGAGACAGTCAGAGGGTATTTATTGAACTAGGTTTGTCTGGTCAACCACCGCGTCCTCAGTGGTGTACGGTACAAACCGCATAAATAGGCGTA<br>TCACGAGGCAGAATTTCAGATAAAAAAAATCCTTAGCTTTCGCTAAGGATGATTTCTGGAATTCGCGGCCGCTTCTAGAGACTAGTGGAAGACATGGAGAGAGACCTG<br>CACCATATGCGGTGAGGGAAGACATGGAGTTGACAGCTAGCTCAGTCCTAGGTACTGTGCTAGCTACTAGAGTCACACAGGAAAGTACTAAATGATGATGGTGAGCA<br>AGGGCGAGGCAGTGATCAAGGAGTTCATGCGGTTTAAGGTGCACATGGAGGGCTCCATGAACGGCCACGAGTTCGAGATCGAGGGCGAGGGCGAGGGCCGCCC<br>CTACGAGGGCACCCAGACCGCCAAGCTGAAGGTGACCAAGGGTGGCCCCCTGCCCTTCTCCTGGGACATCCTGTCCCCTCAGTTCATGTACGGCTCCAGGGCCTT<br>CACCAAGCACCCCGCCGACATCCCCGACTACTATAAGCAGTCCTTCCCCGAGGGCTTCAAGTGGGAGCGCGTGATGAACTTCGAGGACGGCGGCGCCGTGACCGT<br>GACCCAGGACACCTCCCTGGAGGACGGCACCCTGATTTACAAGGTGAAGCTCCGCGGCACCAACTTCCCTCCTGACGGCCCCGTAATGCAGAAGAAAACAATGGGC<br>TGGGAAGCGTCCACCGAGCGGTTGTACCCCGAGGACGGCGTGCTGAAGGGCGACATTAAGATGGCCCTGCGCCTGAAGGACGGCGGCCGCTACCTGGCGGACT<br>TCAAGACCACCTACAAGGCCAAGAAGCCCGTGCAGATGCCCGGCGCCTACAACGTCGACCGCAAGTTGGACATCACCTCCCACAACGAGGACTACACCGTGGTGGA<br>ACAGTACGAACGCTCCGAGGGCCGCCACTCCACCGGCGGCATGGACGAGCTGTACAAGAGGTAGTAAAGGTCCAGGCATCAAATAAAACGAAAGGCTCAGTCGAAA<br>GACTGGGCCTTTCGTTTTATCTGTTGTTTGTCGGTGAACGCTCTCTACTAGAGTCACACTGGCTCACCTTCGGGTGGGCCTTTCTGCGTTTATAGCTTATGTCGTTG<br>CGCTCACTGCCCGCTTTCCAGTCGGGAAACCTGTCGTGCCAGCTGCATTAATGAATCGGCCAACGCGCGGGGGTCTCTGCTTATGTCTTCTACTAGTAGCGGCCGC<br>TGCTCCGGCAAAAAAGGGCAAGGTGTCACCACCCTGCCCTTTTTCTTTAAAACCGAAAAGAATCGGTGTGAAAAGTCAGTATCCAGTCGTGTAGTTCTTATTACCT |
| PJV1160 | ATCGGTGTGAAAAGTCAGTATCCAGTCGTGTAGTTCTTATTACCTTTACTTCGCGTTATGCAGGCTTCCTCGCTCACTGACTCGCTGCGCTCGGTCGTTCGGCTGCG<br>GCGAGCGGTATCAGCTCACTCAAAGGCGGTAAGTCGCGTGTTATACGCCCGTTATCCATGGGTATGGACAGTTTTCCCTTTGATATGTAACGCACGTTGTGTCTCAA<br>AATCTCTGATGTTACATTGCACAAGATAAAAATATATCATCATGAACAATAAAACTGTCTGCTTACATAAACAGTAATACAAGGGGTGTTATGAGCCATATTCAACGGGAA<br>ACGTCTTGCTCCCGTCCGCGCTTAAACTCCAACATGGACGCTGATTTATATGGGTATAAATGGGCTCGCGATAATGTCGGGCAAAGTGGTGCGACAATCTATCGCTT<br>GTATGGGAAGCCCGATGCGCCAGAGTTGTTTCTGAAACATGGCAAAGGTAGCGTTGCCAATGATGTTACAGATGAGATGGTCCGTCTCAACTGGCTGACGGAGTTT<br>ATGCCTCTCCCGACCATCAAGCATTTTATCCGTACTCCTGATGATGCGTGGTTACTCACCACCGCGATTCCTGGGAAAACAGCCTTCCAGGTATTAGAAGAATATCCT<br>GATAGTGGTGAAAATATTGTTGATGCGCTGGCCGTGTTCCTGCGCCGGTTACATTCTATTCCTGTTTGTAATTGTCCTTTTAACAGCGATCGTGTATTTCGTCTTGCT<br>CAGGCGCAAAGTCGCATGAATAACGGTTTGGTTGATGCGAGTGATTTTGATGACGAGCGTAATGGCTGGCCTGTTGAACAAGTCTGGAAAGAAATGCACAAGCTCTT<br>GCCATTCTCTCCGGATAGTGTCGTCACTCATGGTGATTTCAGCCTTGATAACCTTATTTTTGACGAGGGGAAATTAATAGGTTGTATTGATGTTGGACGGGTCGGAAT<br>CGCAGACCGTTACCAGGACCTTGCCATTCTTTGGAACTGCCTCGGTGAGTTTTCTCCTAGTTTACAGAAACGGCTTTTTCAAAAATATGGTATTGATAATCCTGATATG<br>AATAAATTGCAGTTTCATTTGATGCTCGATGAGTTTTTCTAATAATACTAGCAGAAATCATCCTTAGCGAAAGCTAAGGATTTTTTTTATCTGATTACCGCCTTTGAGTGA<br>GCGTCGACCTAGTCCTAGCCAGGATACCGTAAAAAGGCCGCGTTGCTGGCGTTATCACGGCACTACACTCGTTGCTTTATCGGTATTGTTATTACAGA |
| PJV1226 | ATCACGGCACTACACTCGTTGCTTTATCGGTATTGTTATTACAGATTTCCATAGGCTCCGCCCCCCTGACGAGCATCACAAAAATCGACGCTCAAGTCAGAGGTGGCG<br>AAACCCGACAGGACTATAAAGATACCAGGCGTTTCCCCCTGGAAGCTCCCTCGTGCGCTCTCCTGTTCCGACCCTGCCGCTTACCGGATACCTGTCCGCCTTTCTCC<br>CTTCGGGAAGCGTGGCGCTTTCTCATAGCTCACGCTGTAGGTATCTCAGTTCGGTGTAGGTCGTTCGCTCCAAGCTGGGCTGTGTGCACGAACCCCCCGTTCAGC<br>CCGACCGCTGCGCCTTATCCGGTAACTATCGTCTTGAGTCCAACCCGGTAAGACACGACTTATCGCCACTGGCAGCAGCCACTGGTAACAGGATTAGCAGAGCGAG<br>GTATGTAGGCGGTGCTACAGAGTTCTTGAAGTGGTGGCCTAACTACGGCTACACTAGAAGAACAGTATTTGGTATCTGCGCTCTGCTGAAGCCAGTTACCTTCGGAA<br>AAAGAGTTGGTAGCTCTTGATCCGGCAAACAAACCACCGCTGGTAGCGGTGGTTTTTTTGTTTGCAAGCAGCAGATTACGCGCAGAAAAAAAGGATCTCAAGAAGATC<br>CTTTGATCTTTTCTACGGGGTCTGACGCACTTCCATCGCTTCTCTTGGGACGTTTCTTAACTGCTGCAAGACTACTTGAACATTTCGACTTATAATCTATCCGATACGC<br>TAGTTGGGCTGCTCAGACTACCAGTCCCGTTCATTGCCATCCCCGTCCAATTTAAGAACGTGTATGATATGCGAAGCACCTGAAATACCTAGTACCTAATACTCCTACA<br>GCCCCTTTATTGTCTTACCGTATTCCGACCTCTACACCGAGGGGTACGTTAATTCGCCAAAACGATCTTGAGTGGCAATATGCCAACTTGAATACCGTGCCACCGTTA<br>CATCGGCCGGCTTGTCGACGACGGCGTCCTCCGTCGTCAGGATCATCCGGGCACCCACGACTATTGACTGCTCTGAGAAAGTTGATTGTTACGATTA |
| PJV1329 | ACCCACGACTATTGACTGCTCTGAGAAAGTTGATTGTTACGATTACATTGCTGGTGGTACAAGGAGACATGTGACTTTCGTTACCCTCGCGTCAAAAAGAGTTTTTAC<br>GAAAGGAAGCATAAGTGACCTGGGACGATCACAAGAAGAATTTTGCTCGCCTGGCGCGAGATGGTGGTTACACCATCGCACAGTATGCCGCCGAGTTTAATCTTAAC<br>CCTAATACCGCACGTCGTTATCTCCGTGCCTTCAAAGAAGACACCAGGACTACGGACAGCCGCAAGCCAAATAAGCCAGTCAGGAAGCCACTAAAAAGCATGATCATT<br>GATCACTCTAATGATCAACATGCAGGTGATCACATTGCGGCTGAAATAGCGGAAAAACAAAGAGTTAATGCCGTTGTCAGTGCCGCAGTCGAGAATGCGAAGCGCCA<br>AAATAAGCGCATAAATGATCGTTCTGATGATCATGACGTGATCACCCGCGCCCACCGGACCTTACGTGATCGCCTGGAACGCGACACCCTGGATGATGATGGTGAAC<br>GCTTTGAATTCGAAGTTGGCGATTACCTGATAGATAACGTTGAAGCGCGGAAGGCCGCGCGCTATGTTGCGTCGGTCCGGGGCCGATGTTCTGGAAACCACTC<br>TTCTGGAAAAGTCTCTTTCTCATCTCCTTATGCTGGAGAACGCCAGGGATACGTGTATTCGCCTGGTGCAGGAAATGCGCGATCAGCAAAAGACGATGATGAAGGT<br>ACTCCGCCTGAATACCGTATCGCGAGCATGCTAAACAGCTGTTCCGCGCAGATAAGCAGCCTGATCAACACCATTTACAGCATCCGGAATAACTATCGAAAAGAAAGC<br>CGGGAGGCGGAAAAGCACGCTTTATCTATGGGGCAAGCTGGCATTGTTAAGCTGGCATACGAACGAAAGCGTGAAAATAACTGGAGCGTGCTGGAAGCGGCTGAAT<br>TCATCGAGGCGCATGGAGGAAAAGTGCCGCCCCTGATGCTGGAGCAAATCAAAGCCGATCTGCGTGCTCCTAAGACCAATACCGATGATGAGGAAAACCAAACAGCA<br>TCTGGCGCTCCATCCCTTGAAGATCTGGATAAAATCGCGCGAGAACGGGCCGCCAGCCGCCGCGCTGATGCCGCATTGTGGATTGAGCATCGTAGAGAAGAAATTG<br>CCGATATCGTCGATACAGGTGGTTATGGTGATGTCGATGCGGAAGGCATAAGTAACGAAGCATGGCTTGAACAGGATCTGGACGAAGACGAGGAGGAAGACGAAGA<br>AGTTACCCGCAAACTGTACGGGGATGATGATTAATGGCCAGAAGTTGCGTAACGGACCCACGTTGGCGCGAGCTTGTGGCGCTATATCGTTATGACTGGATTGCGG<br>CCGCTGATGTGTTGTTTGGGAAGACACCAACCTGGCAGCAGGATGAGATCATTGAGTCCACGCAGCAGGACGGCAGTTGGACAAGTGTGACCTCCGGCCATGGTA<br>CTGGTAAATCTGATATGACGAGTATCATTGCAATACTCTTCATCATGTTTTTCCCCGGCGCTCGCGTCATTCTGGTCGCTAACAAAAGACAGCAAGTCCTTGATGGTAT<br>TTTCAAATACATAAAGAGCAATTGGGCTACTGCTGTTAGCAGATTCCCGTGGTTGTCCAAGTATTTCATTCTTACAGAAACGTCTTTTTTTGAGGTGACTGGCAAGGGT<br>GTTTGGACAATATTGATAAAGTCCTGTCGTCCCGGAAATGAGGAGGCGTTGGCTGGTGAACACGCCGATCATCTCTTGTATATCATCGACGAAGCGAGCGGTGTGA<br>GTGATAAAGCATTCAGTGTGATAACAGGTGCGCTGACCGGTAAGGATAACCGTATTCTGCTTCTTTCCCAGCCTACGCGACCTTCTGGCTATTTCTACGATAGTCACC<br>ACAGACTAGCTATTCGCCCGGGAAATCCTGATGGATTGTTTACTGCGATAATACTGAATAGTGAAGAATCTCCGCTTGTAGATGCAAAATTTATACGAGCAAAACTTGC<br>GGAGTATGGCGGTCGTGATAACCCCATGTACATGATCAAAGTACGTGGTGAATTTCCCAAATCTCAAGATGGCTTTCTTCTTGGTCGTGATGAGGTTGAGCGGGCGA<br>CGCGGCGAAAGGTCAAGATTGCCAAAGGATGGGGCTGGGTTGCATGTGTTGACGTTGCTGGTGGCACAGGACGAGATAAGTCCGTTATTAATATCATGATGGTGTC<br>CGGCCAGCGAAATAAACGCCGTGTAATCAACTATCGTATGCTGGAATACACAGACGTTACAGAAACGCAGTTAGCCGCCAAGATTTTCGCAGAATGTAACCCAGAACG<br>GTTCCCGAACATAACCATAGCTATTGATGGCGATGGCTTGGGGAAAGCACGGCTGATCTAATGTACGAACGCTATGGCATTACCGTCCAGCGTATCCGCTGGGGTA<br>AAAAGATGCACAGCCGTGAAGATAAAAGCCTTTATTTCGATATGCGCGCTTTCGCGAATATTCAGGCGGCAGAAGCTGTAAAATCCGGGCGTATGAGGCTTGATAAG<br>GGGGCTGCGACTATAGAGGAAGCATCTAAGATACCGGTAGGGATAAATTCCGCAGGTCAATGGAAGGTGATGAGCAAGGAAGATATGAAGAAAAAACTCAACCTGCA<br>CTCTCCGGACCATTGGGATACATATTGTTTCGCTATGTTGGCGAACTATGTTCCCCAAGATGAAGTGCTTAGCGTCGAAGACGAAGCGCAGGTTGATGAAGCTCTGG<br>CATGGCTTAATGAATAATAGTCCGGCAATTAAAAAAGCGGCTAACCACGCCGCTTTTTTTACGTCTGCACTCGGTACCAAATTCCAGAAAAGAGGCCTCCCGAAAGGG<br>GGGCCTTTTTTCGTTTTGGTCCCTCCTTCGTCCACCAGCAATGTTACTTACGACACTCCGAGACAGTCAGAGGGTATTTATTGAACTA |
| PJV1345 | AAAGGCTATCGCTGTGTCTAAGTATTCGTCTGTTTTGATAGATGACAGTACACGCCCACGTAGATTTCAGATCCACTCTCTGAATCCTGAATTCTTTCGCCGTAGCCG<br>CGCTCCGCGCCGGTTTGATGAAAAACTCCAGAACGAACTCTATATGACGCAGGACGAAAAGGAACGCCGGGAGCACCAGCCTTGGGTAATGGCGCGTCAACTTTTC<br>AATAAGGTGGCCCGTCAGCACCGTCATTACGGTAATGCCACATCCGCACGTATCTGATTGATTGCTTGCCCGTTCCGGGCCTTTTGTCTTTAAAAAGAAACCTCCGCA<br>TTGCGGAGGTTTCGCCTTTTGATACTCTGTCTGAAGTAATTCTTGCCGCAGTGAAAAATGGCGCCCATCGGCGCCATTTTTTTATGCTTCCATTAGAAAGCAAAAAGC<br>CTGCTAGAAAGCAGGCTTTTTTGAATTTGGCTCCTCTGACAGTACTGGCGGGGTTGAACGGTCAGGTATCACTATAATCAAAACCCCTCGATATATGCATTAATCTCG<br>CCGGTCCGATGAACTGACGGTCATGTAGCTAGTATGAGCCGCGAAGAGGGACGGAGACATGCCAAACGCGAACAAGGTCATCCCTATGCCGAACAATATGTCGCCC<br>TGGCTTTATAGCAGATTACGGTCAGCGCCGCTATATACATAAACTGTGACTAACATATTATCCCTGTGTATAACACCAGTTCGATCCGAAACCCAAATGTATCAGAGTCA<br>CCGTTAATCCACTAAAGGTCTGAGTACTTGTGCACCATATGCGGTGAGGGATGATCGGCACGTAAGAGGTTCCAACTTTCACCATAATGAAATAAGATCACTACCGGG<br>CGTATTTTTTGAGTTGTCGAGATTTTCAGGAGCTAAGGAAGCTAAAATGGAGAAAAAAATAACGGGCTACACAACCGTTGATATATCAGCCAGTGGCATAGAAAAGAACAT<br>TTCGAGGCATTTCAGTCTGTTGCTCAATGCACCTACAACCAGACAGTTCAGCTGGATATCACTGCCTTTCTGAAGACCGTAAAGAAAAACAAACACAAGTTTTATCCGG<br>CCTTTATTCACATTCTTGCGCGCCTGATGAACGCCCATCCGGAATTTCGTATGGCAATGAAAGACGGGGAGTTGGTGATTTGGGATTCGGTTCATCCATGTTATACC<br>GTCTTCCACGAACAGACGGAAACTTTCTCCTCTCTGTGGAGTGAATACCACGACGATTTTCGCCAGTTCCTCCACATCTATAGCCAAGATGTGGCGTGCTACGGAGAA<br>AATCTGGCGTATTTCCCTAAAGGGTTTATTGAGAATATGTTTTTTGTTTCGGCCAATCCCTGGGTGTCCTTCACCTCATTCGATTTGAATGTGGCAAATATGGACAACT<br>TTTTTGCTCCGGTATTCACCATGGGTAAATATTATACGCAAGGCGACAAAGTGTTAATGCCGCTGGCCATTCAGGTCCATCATGCGGTGTGTGATGGTTTTCATGTG<br>GGGAGGATGTTAAACGAACTACAACAATACTGCGATGAATGGCAGGGAGGTGCGTGATAAAGGTCCAGGCATCAAATAAAACGAAAGGCTCAGTCGAAAGACTGGGC<br>CTTTCGTTTTATCTGTTGTTTGTCGGTGAACGCTCTCTACTAGAGTCACACTGGCTCACCTTCGGGTGGGCCTTTCTGCGTTTATACTCATTAACCATGCCGGATGGA<br>AACTATTGCGCGCTTTCGGGGTTGTCGTTTACTGGCTGCCCCTTCTTAGTTTTGCGGCTGCGCGTAACTGATGCGGCTGATTTGACCTTTTTCTCTTCGCGAGTGAT<br>GGCAATTTGTTTTTTTACATTTTCAATATCTGCCAGGCGATATATTTTTGCCTGCGGCCAGCGGTCGCAGATGATCGGTTCTATAGAGTCATAAAGGCTAAATTTTGCT<br>TTCTCAAATTCACCGTTGATGATGATTCCATCTCGGAGAGTTTCATCGCAGATAAACACGCA |

**Supplementary Table 8.** Names and sequences of synthesized DNA fragments used in this study.

| Plasmid Name | Part 1 | Part 2 | Part 3 | Part 4 |
|---|---|---|---|---|
| pT7gp17 | PJV1063 | PJV1161 | PJV720 | - |
| pT7kan-mScarlet | PJV684 | PJV686 | PJV1160 | PJV1226 |
| pP1kan-mScarlet | PJV1329 | PJV686 | PJV1160 | PJV1226 |

**Supplementary Table 9. Gibson assembly schemes.** DNA parts in **Supplementary Table 6** were Gibson assembled according to the following schemes, as described in the Methods.

| Name | Sequence |
|---|---|
| JDL001 | CTGTGTCCCTTGTCTCATGAGCGGATAC |
| JDL002 | CACTGTGAGACCTTGTTCATGTGTGTTC |
| PJV001 | TATAACGTTTTTGAACACACATGAACAAGGTCTCACAGTGTACGGACCTAAAGTTCCCCC |
| PJV002 | ATTACGCGATGACAGTAGACAACCTTTCCG |
| PJV003 | TGCAGCAATACCGGAAAGGTTGTCTACTGT |
| PJV004 | ATATGTCTCCTCATAGATGTGCCTATGTGG |
| PJV005 | ACTTGTGACTCCACATAGGCACATCTATGA |
| PJV006 | GAATAACCTGAGGGTCAATACCCTGCTTGT |
| PJV007 | GACATGATGGACAAGCAGGGTATTGACCCT |
| PJV1253 | TTGACCTCCTTAAAGTAAATCTAAGAGACTACAGGGAGAA |
| PJV012 | TTGGTAAATCACAAGGAAAGACGTGTAGTC |
| PJV008 | ACATTCAAATATGTATCCGCTCATGAGACAAGGGACACAGAGAGACACTCAAGGTAACAC |
| PJV1254 | TAGGGAGAGGCGAAATAATCTTCT |
| PJV1255 | ACCTTGTACCTCCTTGAGAGTC |

**Supplementary Table 10. Primers for barcoded T7 genome assembly.** Primers used to amplify sections of the T7 genome that were subsequently assembled in a yeast artificial chromosome and then transformed into *E. coli*, rebooting the 7898-barcoded T7 variant.

| Name | Sequence |
| --- | --- |
| 8673F | GGACGTTTCTTAACTGCTGCA |
| 8673P | 6FAM-ACGCTAGTT+GGGCTGCTCAGA-Iowa_Black_FQ |
| 8673R | GGCAATGAACGGGACTGGTA |
| 7898F | CATCCCTATGCCGAACAATATG |
| 7898P | HEX-AGCAGATTA+CGGTCAGCGCC--Iowa_Black_FQ |
| 7898R | CGGATCGAACTGGTGTTATAC |

**Supplementary Table 11. Primers and Probes used in the optimized ddPCR assay.** + denotes locked nucleic acid.
